# Larval zebrafish display dynamic learning of aversive stimuli in a constant visual surrounding

**DOI:** 10.1101/2021.01.30.428906

**Authors:** Jiale Xu, Romelo Casanave, Su Guo

## Abstract

Balancing exploration and anti-predation are fundamental to the fitness and survival of all animal species from early life stages. How these basic survival instincts drive learning remains poorly understood. Here, employing a light/dark preference paradigm with well-controlled luminance history and constant visual surrounding in larval zebrafish, we analyzed intra- and inter-trial dynamics for two behavioral components, dark avoidance and center avoidance. We uncover that larval zebrafish display short-term learning of dark avoidance with initial sensitization followed by habituation; they also exhibit long-term learning that is sensitive to trial interval length. We further show that such stereotyped learning patterns is stimulus specific, as they are not observed for center avoidance. Finally, we demonstrate at individual levels that long-term learning is under homeostatic control. Together, our work has established a novel paradigm to understand learning, uncovered sequential sensitization and habituation, and demonstrated stimulus specificity, individuality, as well as dynamicity in learning.

## Introduction

Learning while being exposed to a stimulus (i.e., non-associative learning) is of great importance in that it triggers intrinsic constructs for subsequent recognition of that stimulus and provides a foundation for associative learning (e.g., learning about relations between stimuli in Pavlovian conditioning and stimuli responses-outcomes in instrumental conditioning). Non-associative learning precedes associative learning in the evolutionary sequence and involves a broad range of behavioral phenomena including habituation, sensitization, perceptual learning, priming and recognition memory (Ioannou and Anastassiou-Hadjicharalambous, 2018; Pereira and van der Kooy, 2013).

As the simplest learning form, habituation is defined as the progressively reduced ability of a stimulus to elicit a behavioral response over time (Glanzman, 2009; Rankin et al., 2009; Thompson, 2009). Such a response reduction is distinguished from sensory adaptation and motor fatigue and is often considered adaptive in that it helps animals to filter out harmless and irrelevant stimuli (Rankin et al., 2009). Since an early study of EEG arousal in cats (Sharpless and Jasper, 1956), the habituation phenomenon has been widely documented in invertebrates such as *C. elegans* (Ardiel et al., 2016; Giles and Rankin, 2009; Rankin and Broster, 1992; Rose and Rankin, 2001) and *Aplysia* (Glanzman, 2009) as well as in vertebrates such as rodents (Arbuckle et al., 2015; Bolivar, 2009; Salomons et al., 2010), zebrafish (Best et al., 2008; Pantoja et al., 2020; Randlett et al., 2019; Roberts et al., 2016; Wolman et al., 2011) and humans (Bornstein et al., 1988; Coppola et al., 2013).

Accompanying habituation is a process termed sensitization, which by contrast enhances responses to a stimulus over time (Kalivas and Stewart, 1991; McSweeney and Murphy, 2009; Robinson and Becker, 1986). This counterpart of habituation may also be adaptive if it helps animals avoid potentially risky or costly situations (Blumstein, 2016; King et al., 2007). Like habituation, sensitization has also been documented in a phylogenetically diverse set of organisms (Cai et al., 2012; Kirshenbaum et al., 2019; Tran and Gerlai, 2014; Watkins et al., 2010), suggesting an evolutionarily conserved biological underpinning for both processes. Furthermore, these simple learning forms are observed in various functional outputs of nervous systems ranging from simple reflexes (Blanch et al., 2014; Pantoja et al., 2020; Pinsker et al., 1970; Randlett et al., 2019) to complex cognitive phenotypes (Bolivar, 2009; Leussis and Bolivar, 2006; Thompson and Spencer, 1966) and may represent deeper neurobiological constructs associated with anxiety-memory interplay (Morgan and LeDoux, 1995; Ruehle et al., 2012; Sullivan and Gratton, 2002). Therefore, understanding basic building blocks of habituation and sensitization is essential to fully understand complex behaviors.

Habituation and sensitization have been reported with short-term (intrasession) and long-term (intersession) mechanisms in a number of systems (Rankin et al., 2009; Thompson, 2009). Short-term mechanisms sensitize or habituate a response within a session (Byrne and Hawkins, 2015; Leussis and Bolivar, 2006; Meincke et al., 2004; Rahn et al., 2013). In contrast, long-term mechanisms retain memory formed in previous session and use it to modify behavioral responses in a subsequent session (Rankin et al., 2009).

Although both short- and long-term learning and memory have been demonstrated in young larval zebrafish when exposed to a single stimulus (O’Neale et al., 2014; Randlett et al., 2019; Roberts et al., 2016; Wolman et al., 2011), so far, most paradigms use unnatural stimuli and are designed without integrating sensitization and habituation in the same paradigm. The latter limitation is crucial as the influential dual-process theory, proposed by Groves and Thompson in 1970, suggests that the two learning forms interact to yield final behavioral outcomes and therefore assessment of only one process might be confounded by alteration in the other process (Meincke et al., 2004).

In this study, we examined stimulus learning in a large population of larval zebrafish using a light/dark preference paradigm over four trials across two days. Light/dark preference as a motivated behavior is observed across the animal kingdom (Bourin and Hascoët, 2003; Gong et al., 2010; Lau et al., 2011; Serra et al., 1999). Larval zebrafish display distinct motor behaviors that are sensitive to the intensity of both pre-adapted and current photic stimuli (Burgess et al., 2010; Burgess and Granato, 2007; Facciol et al., 2019). In our paradigm with well-controlled luminance history and constant visual surrounding, larval zebrafish generally display dark avoidance and center avoidance (also known as thigmotaxis) with heritable individual variability and are considered fear or anxiety-related (Bai et al., 2016; Dahlén et al., 2019; Schnörr et al., 2012; Steenbergen et al., 2011; Wagle et al., 2017). From an ethological perspective, the extent of avoidance is likely a readout of the circuitry that balances anti-predation (i.e., avoid the dark and the center) and free exploration (i.e., approach the dark and the center). As described in the Results section below, we have uncovered stimulus-specific temporal dynamicity of learning (both short-term and long-term), as well as individual differences in learning that are under homeostatic control.

## Results

### The dark avoidance behavior displays intra-trial sensitization followed by habituation

We developed a population of 1680 healthy wild-type EK larval zebrafish. Each individual was pre-adapted to a well-controlled luminance background, then introduced into a half-light and half-dark choice chamber and video recorded in four trials of 8-minute each. During each trial, animals were exposed to a constant visual surrounding and had the freedom to navigate the entire arena, thereby mimicking possible encounters in nature. The inter-trial time interval was ~2 hours between trial 1-2 and 3-4, whereas the inter-trial interval was ~22 hours between trial 2-3. Two behavioral responses were analyzed: the dark avoidance behavior was measured by the light/dark choice index (LD-CI) and the center avoidance behavior was quantified by the periphery/center index (PC-CI) (**Figure 1**). Previous studies have shown that while individual variability exists, larval zebrafish generally find the dark more aversive than the light environment, and the center more aversive than the periphery of the arena. These preferences likely reflect anti-predatory responses (Bai et al., 2016; Dahlén et al., 2019; Prut and Belzung, 2003; Schnörr et al., 2012; Treit and Fundytus, 1988; Wagle et al., 2017).

**Figure 1.**
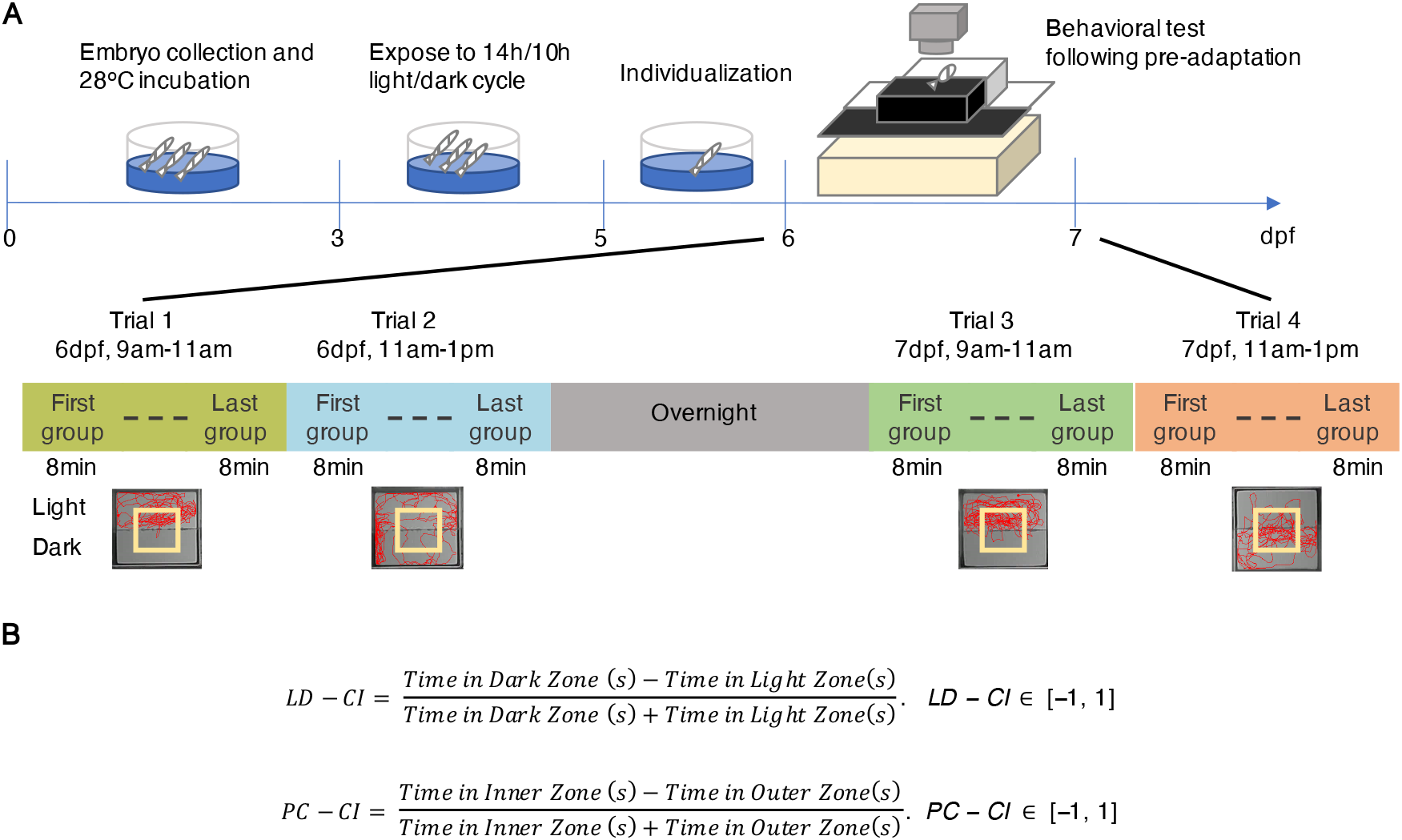
Schematic design of the behavioral assay for characterizing stimulus learning in larval zebrafish. (**A**). Larvae were tested in a chamber (4cm (L) × 4 cm (W) × 1.5 cm (H)) with a physical light-dark division (top panel) and a virtual center-periphery division (indicated by the 1cm × 1cm yellow square border in the bottom panel). Individual behavior was recorded in four 8-min trials scheduled from 6dpf to 7dpf (middle panel). A group of sixteen individuals were recorded at a time for an 8-min trial and up to 192 individuals were tested in a 2hr session (middle panel). The early trials on each day was performed from 9am to 11am followed by the later trials from 11am to 1pm. Experiments were weekly repeated to test a total of 1680 larvae and the movement recordings were subsequently digitized to compute the time spent in the divided zones (bottom panel). (**B**) Equations for quantifying the light-dark choice index (LD-CI) and the periphery-center choice index (PC-CI).

Both types of behavioral responses could potentially be subjected to non-associative learning, resulting in short- and long-term memories that modify the original responsiveness to aversive stimuli. We first assessed short-term learning by characterizing behavioral dynamics within a trial. We profiled each trial with sixteen 30-second time bins and took the population (n=1680) mean of LD-CI and PC-CI for each bin to uncover intra-trial dynamics of the two behaviors respectively. We found a decline of LD-CI in early time bins followed by a progressive increase throughout the rest of time bins (**Figure 2A**). The early decline of LD-CI indicated an increase of dark avoidance (i.e., sensitization), whereas the later increase of LD-CI demonstrated a decrease of dark avoidance (i.e., habituation). Furthermore, increased dark avoidance in the sensitization period coincided with an increase of swim velocity that was stabilized during the subsequent habituation period (**Figure S1**). Together, these results demonstrate that larval zebrafish display short-term learning of dark avoidance by an initial sensitization followed by habituation.

**Figure 2.**
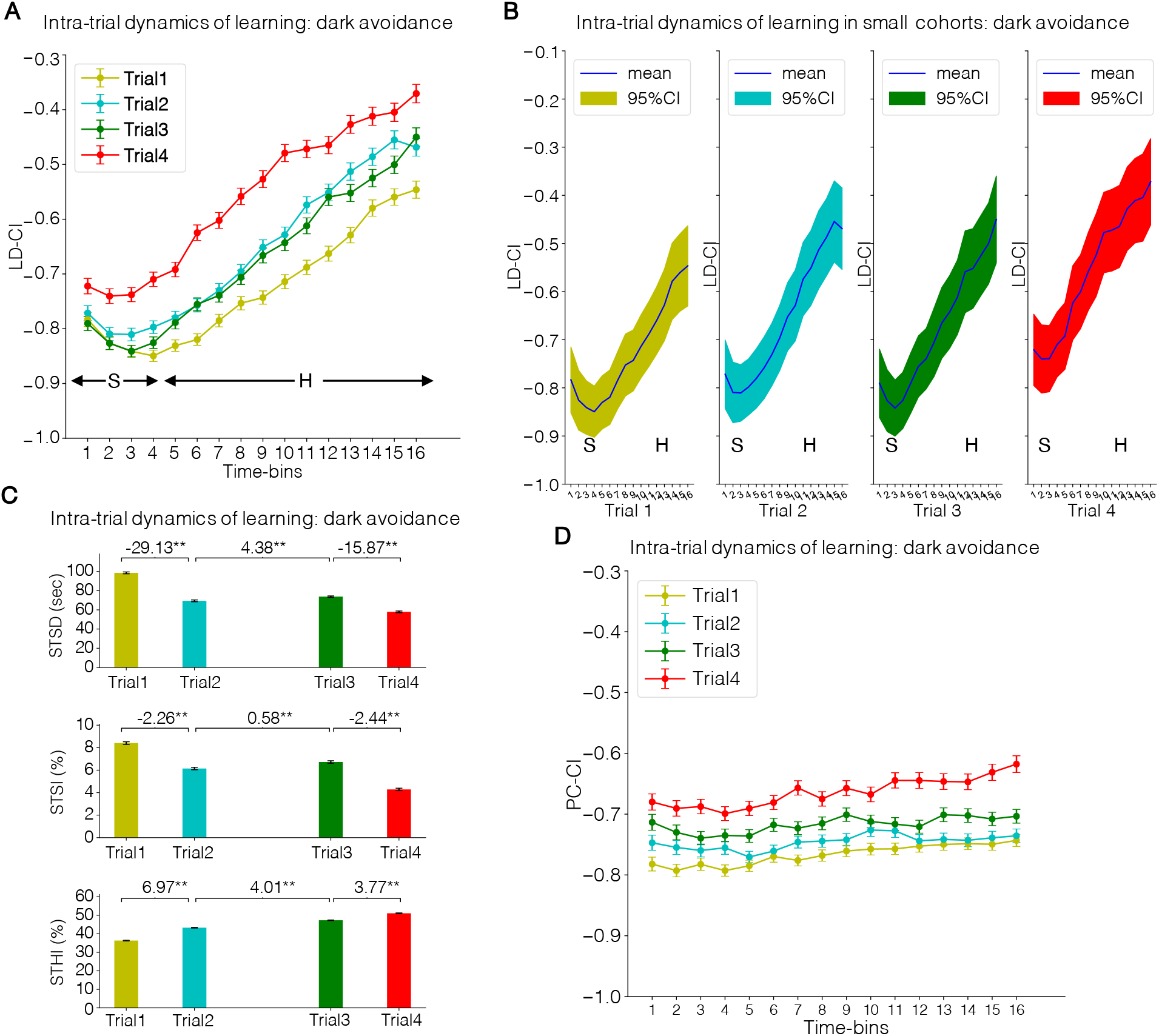
Larval zebrafish display dynamic short-term learning of dark avoidance with initial sensitization followed by habituation. (**A**) Intra-trial kinetics of LD-CI. In all four trials, LD-CI was initially decreased during the first several bins followed by a continuous increase over the rest bins, indicating that dark avoidance was first sensitized for a short period before it was habituated. (**B**) A collection of intra-trial kinetics of LD-CI in small cohorts of 200 individuals. A total of 1000 iterations of random subsampling demonstrated the sequential occurrence of sensitization and habituation is a characteristic detectable in small cohorts of 200 individuals. (**C**) Quantitative characterization of intra-trial kinetics. Three parameters were computed (see methods) to quantify intra-trial kinetics of LD-CI. Comparisons between trials were performed with vectors of each parameter (paired one-tail t-test, \*\**p*<0.01, \**p*<0.05) (**D**) Intra-trial kinetics of PC-CI. Compared to LD-CI, changes of PC-CI was observed with a much lower magnitude, suggesting a minimal effect of habituation on aversive response to the arena center. H, habituation; S, sensitization; STSD, short-term sensitization duration; STSI, short-term sensitization index; STHI, short-term habituation index. LD-CI, light/dark choice index; PC-CI, periphery-center choice index. A paired one-tailed t-test is used for panel C. For A and D, n=1680 individuals. For B and C, n=1000 iterations. Each iteration sampled 200 individuals.

To determine whether this phenomenon is observable in smaller cohorts or whether it is only detectable in the large cohort of 1680 individuals, we performed 1000 iterations of random subsampling of 200 individuals and calculated the sub-population (n=200) mean of LD-CI for each 30-sec time bin (see Methods). Similar to the entire population, most of the 1000 small cohorts underwent sensitization in early time bins before they transitioned to habituation (**Figure 2B**). This analysis reinforces that larval zebrafish learn dark stimulus through an initial short sensitization followed by a subsequent habituation, which can be consistently detected in cohorts as small as 200 individuals.

To further characterize the intra-trial dynamic learning of dark avoidance, we calculated three parameters for every subsampled population: 1) short-term sensitization duration (STSD), 2) short-term sensitization index (STSI), and 3) short-term habituation index (STHI) (**Figure 2C**). These analyses uncovered that Trial 1 had the longest STSD (99-sec), which also resulted in the highest STSI (9.7%) and the lowest STHI (36.3%) (**Figure 2C**). Across the four trials, cohorts displayed a decrease of both STSD and STSI upon a stimulus re-exposure after two hours (between trial 1-2 and trial 3-4), while a retest after an overnight break (between trial 2-3) resulted in an increase of both parameters. In contrast, the STHI showed steady increase across the four trials.

Intriguingly, the dynamic learning of dark avoidance (i.e., sensitization followed by habituation) was not observed with regard to the center avoidance behavior, which underwent a marginal change within the trial, suggesting minimal short-term learning of the periphery-center stimulus (**Figure 2D**). Taken together, larval zebrafish display dynamic short-term learning in a stimulus-specific manner: when faced with the light/dark stimulus, larval zebrafish display initial sensitization followed by habituation.

### The dark avoidance behavior displays long-term habituation or sensitization in an inter-trial interval-dependent manner

Long-term habituation, which lasts from hours to days, was measured by inter-trial comparisons of behavioral indices. While inter-trial dynamics could be glimpsed from data presented in **Figure 2A** and **2C**, we further calculated average choice indices for each of the four trials and compared them between consecutive trials. As shown in **Figure 3A**, larvae showed less dark avoidance in a second trial (Trial 2, LD-CI=−0.65±0.009) conducted two hours after the first trial (Trial 1, LD-CI=−0.72±0.008), suggesting habituation. A similar long-term habituation phenomenon was also observed when comparing trial 4 to 3 at 7dpf (Trial 3, LD-CI=-0.67±0.008; Trial 4, LD-CI=-0.56±0.009;). However, when comparing the two trials with an overnight interval (i.e. Trial 3 to 2), we uncovered a slight but significant increase of dark avoidance, suggesting sensitization. These results indicate that an overnight time interval has not only erased the habituation memory but also further sensitized the dark avoidance behavioral response. Intriguingly, such a time-dependent learning outcome closely mirrored the observed dynamicity in STSD and STSI (see **Figure 2C**), suggesting that the long-term effects of stimulus learning appear to abide by short-term learning performance.

**Figure 3.**
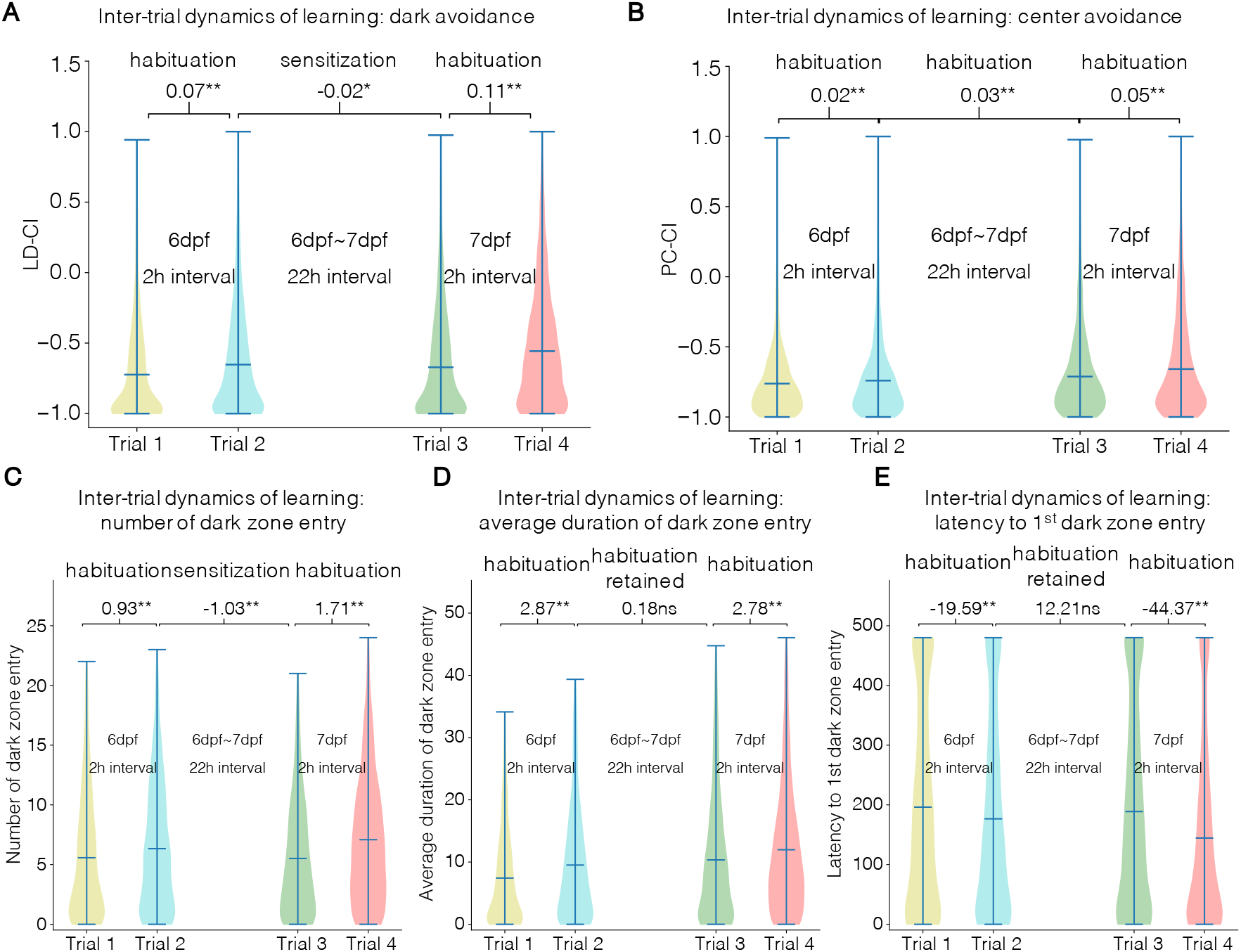
Long-term stimulus learning exhibit different effects depending on inter-trial interval. (**A**) Population mean of LD-CI significantly increased over two successive trials performed with a 2h interval, indicative of habituation (Trial 2 vs Trial 1, mean LD-CI diff. = 0.07, \*\**p*<0.01; Trial 4 vs Trial 3, mean LD-CI diff. = 0.11, \*\**p*<0.01). A decrease of mean LD-CI was detected across the two trials performed with an overnight interval, indicative of sensitization (Trial 3 vs Trial 2, mean LD-CI diff. = −0.02, \**p*<0.05). (**B**) Population mean of PC-CI were observed with slight yet still distinguishable increments for the two 2h intervals (Trial2 vs Trial 1, mean PC-CI diff. = 0.02, \*\**p*<0.01; Trial 4 vs Trial 3, mean PC-CI diff. = 0.05, \*\**p*<0.01) as well as the overnight interval, indicative of habituation (Trial 3 vs Trial 2, mean PC-CI diff. = 0.03, \*\**p*<0.01). (**C-E**) Two components positively correlated with LD-CI, the number of dark zone entry (**C)** and average duration of dark entry (**D**), were increased significantly over the 2h intervals (Trial 2 vs Trial 1, mean diff. of number of dark zone entry = 0.93, \*\**p*<0.01, mean diff. of average duration of dark zone entry = 2.87, \*\**p*<0.01; Trial 4 vs Trial 3, mean diff. of number of dark zone entry = 1.71, \*\**p*<0.01; mean diff. of average duration of dark zone entry = 2.78, \*\**p*<0.01). Like the LD-CI, The population also demonstrated a detectable overnight decrease in its number of dark zone entry (Trial 3 vs Trial 2, mean diff. = −1.03, \*\**p*<0.01) while the average duration of dark entry was not detected with a change of significance (Trial 3 vs Trial 2, mean diff. = 0.18, ns *p* >0.05). As a negatively correlated component, latency to the first dark zone entry (**E**) was significantly decreased over the 2h intervals but its overnight increase was not statistically detectable (Trial 2 vs Trial 1, mean diff. = −19.59, \*\**p*<0.01; Trial 4 vs Trial 3, mean diff. = −44.37, \*\**p*<0.01; Trial 3 vs Trial 2, mean diff. = 12.21, ns *p* >0.05). A paired one-tailed t-test is used for all panels with n=1680 individuals.

In contrast to dark avoidance, long-term habituation of center avoidance steadily progressed across all four trials (**Figure 3B**), with a magnitude that is however smaller than that of dark avoidance. Given the intriguing patterns of habituation and sensitization of dark avoidance behavior, in subsequent sections, we will focus on this behavior; wherever applicable, we will also analyze center avoidance as a comparison.

To verify whether the observed behavioral differences between trials are truly due to learning rather than a simple difference in test timing, we compared the first groups and last groups of Trial 1 that were tested ~2 hours apart (**Figure 1A**). No significant differences in their choice indices were detected (**Figure S2**), indicating that the observed behavioral differences between trials are a result of previous experience.

### Number of dark zone entry undergoes long-term habituation or sensitization in an inter-trial interval dependent manner

The complex trait of dark avoidance could be dissected into several sub-components. Accordingly, the dynamics of dark avoidance across the four trials could be explained by plasticity in one single sub-component or in multiple sub-components. To further investigate the learning rules of dark avoidance, we quantified inter-trial changes of three sub-components: 1) the number of dark zone entry, 2) average duration of dark zone entry, and 3) the latency to the first dark zone entry.

Our analysis uncovered that, while long-term habituation was observable for all three sub-components, only the first sub-component “*the number of dark zone entry*” showed a significant overnight sensitization (**Figure 3C-E**). These results suggest that all three sub-components develop habituation with a possibly overlapping mechanism. The fact that the sensitization only occurs in one of the three sub-components suggests that larval zebrafish can retain long-term (24 hours) habituation memory (in the case of second and third sub-components) but can also replace it with sensitization (in the case of first sub-component), which might be adaptive if it helps the animals avoid potentially risky or costly situations.

### Stimulus learning occurs with individual variability

While stimulus learning displayed stereotypical patterns as we have shown at the population and subpopulation levels, we also observed a considerable spread in distribution across individuals. To further explore the underlying principles, we developed a long-term learning index (LTLI) that quantifies behavioral changes at individual levels between two successive trials (**Figure 4A, Equation 2**). A value of LTLI greater than 1 indicates habituation while less than 1 implies sensitization of that behavior. Individuals with the minimum LD-CI value of −1 in either or both trials cannot be properly characterized with the LTLI equation and were therefore excluded for plotting the distribution curve (see below). Nevertheless, they were still classifiable as habituated or sensitized individuals depending on their behavioral differences between successive trials. Of note, individuals with the minimum LD-CI value of −1 in successive trials were separately categorized as non-learners.

**Figure 4.**
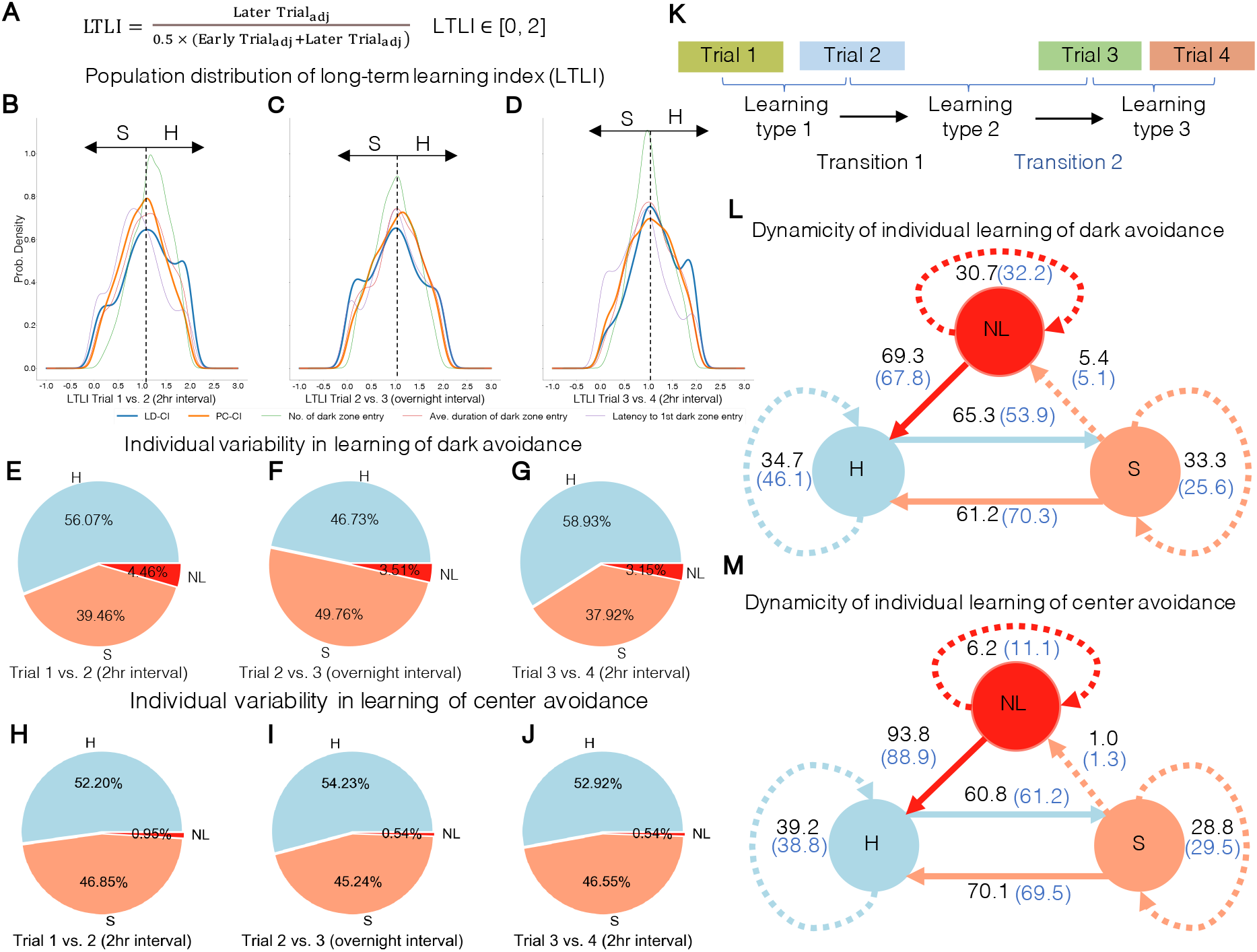
Long-term stimulus learning occurs with individual variability and under homeostatic control. **(A-D)** Long-term learning index (LTLI) computed at individual level unraveled learning performance with individuality. (**E-G**) Based on the LTLI of LD-CI, the entire population was divided into three groups corresponding to three distinct learning types of dark avoidance: habituation (or H, blue), sensitization (or S, orange) and non-learning (or NL, red). With the first learning process at 6dpf (**E**), more individuals display a habituation learning (56.07%) as opposed to sensitization learning (39.46%). A similar but more biased situation was observed in the third learning process at 7dpf (**G**). In contrast, the second learning process was detected with a reversed scenario (**F**) where learning was biased towards sensitization (49.76%) as opposed to habituation (46.73%). In each of the three learning processes, about 3% to 4% of the population were detected as a non-learner. (**H-J**) Using the same criteria, the population were divided into three groups that differently learn center avoidance. Unlike the learning of dark avoidance, the portions of the sensitized and habituated individuals remained steady with the non-learning behavior detected in less than 1% of the population. (**K-M**) Three learning types were assigned to an individual based on its differences of LD-CI in two successive trials (**K**). A transition diagram was constructed for each avoidance behavior to illustrate individual learning dynamicity. Each of the three possible preceding learning types are represented by a circle filled with the color corresponding to the pie chart. Conditioned on a certain preceding learning type, a solid arrow is used to indicate the major transition while other minor transitions were symbolized by dashed arrows. Transition probabilities were designated nearby each arrow for the first (black) and second transition (blue). Throughout all four trials, a preceding learning type is more likely to be transitioned to the opposite learning type than to be maintained as the same. This tendency applies for learning of both dark (**L**) and center (**M**) avoidance.

Although the population mean indicates a pervasive long-term habituation or sensitization depending on trial interval lengths (**Figure 3**), the spread distribution curve of LTLI uncovered individuality in stimulus learning (**Figure 4B-D**). The LTLI distribution of dark avoidance deviates from a gaussian distribution by displaying a higher frequency with values close to both extremes (i.e., “shoulders” observed in the blue graphs of **Figures 4B-D**). This represented the individuals that displayed a LD-CI close to the lower limit (−1) in one of the two consecutive trials (indicating an immediate withdrawal from the dark zone after entry).

In the trials separated by a 2h interval, we found more than 56% of the population underwent habituation learning at both 6dpf and 7dpf compared to a less than 40% observed with sensitization learning (**Figure 4E, 4G)**. In contrast to 2h intervals, re-exposure after an overnight interval resulted in a reversed scenario where more individuals showed sensitization (49.76%) as opposed to habituation (46.73%) (**Figure 4F**). A small portion (3.15% ~4.46%) of non-learners were always detected throughout all trials.

Unlike the deviated learning distribution of dark avoidance, learning of center avoidance showed a near normal distribution throughout all four trials with a slight bias towards the habituation side (orange graphs, **Figure 4B-D**). Percent of individuals that displayed habituation or sensitization learning remained steady across the trials (**Figure 4H-J**). In addition, less than 1% of the population were detected as nonlearners. These observed differences further support the notion that learning of the two avoidance behaviors are governed by different principles.

### Individual learning is under a homeostatic control

Next, we wondered how an individual’s learning performance varies across trials. In other words, will a habituated individual defined in Trial 1-2 continue the same type of learning in subsequent trials (Trial 2-3, and Trial 3-4)? To address this question, we constructed a transition diagram that illustrates an individual’s probability of displaying a certain learning type following its previous learning type. Two transitions of learning performance can be defined in our four-trial paradigm: First transition compares learning that occurred during Trial 1-2 vs. Trial 2-3, and second transition compares learning that occurred during Trial 2-3 vs. Trial 3-4 (**Figure 4K**). For the first transition, a majority of previously habituated individuals (65.3%) were transitioned into a sensitization learning (**Figure 4L**). Likewise, a majority of previously sensitized individuals (61.2%) were transitioned to habituation learning behavior. A similar trend was observed for the second transition: A preceding habituation drove a preference for subsequent sensitization (53.9%), and a preceding sensitization was followed by preferential habituation (70.3%). In addition, non-learners were transitioned into habituation learners in the following trial with a probability (67.8%-69.3%) about twice as high as those remaining as a non-learner (30.7%-32.2%).

The fact that learning behavior tends to switch to the opposite type as opposed to maintaining the same type was also observed with respect to the center avoidance behavior (**Figure 4M**). Together, such an alternation between different learning patterns suggest an underlying homeostatic process that keeps the extent of avoidance behavior within a steady range for optimal adaptation to environmental changes.

While a majority of individuals displayed homeostatically balanced learning patterns, it was worth noting that a small percentage of individuals did not. For instance, across the four trials, an estimated ~3% individuals (39.46% × 33.3% × 25.6%) continued to become sensitized toward the dark stimulus, whereas ~9% individuals (56.07% × 34.7% × 46.1%) continued to become habituated toward the dark stimulus. Thus, these rarer individuals likely represent opposite sides of the spectrum underlying individual learning patterns.

### Long-term learning is more apparent with increased sample size

In an effort to assess the importance of sample size for identifying long-term learning patterns, we examined the power for detecting inter-trial differences in dark avoidance behavior (LD-CI) in sub-samples of the full data set (**Figure 5**). For detecting long-term habituation (i.e., LD-CI changes between trial 1-2 and trial 3-4), the power increased exponentially from 0.5 with a sample size of 100 to the maximum of 1.0 with a sample size of 500. As for the overnight sensitization, the power increases in a linear manner from 0.22 with 100 larvae to 0.65 with 1600 larvae. Our results suggest a necessity of employing a sufficient sample size in order to detect long-term learning patterns, as a larger sample not only minimizes the standard error but also more closely approximate the entire population (Everitt and Skrondal, 2002).

**Figure 5.**
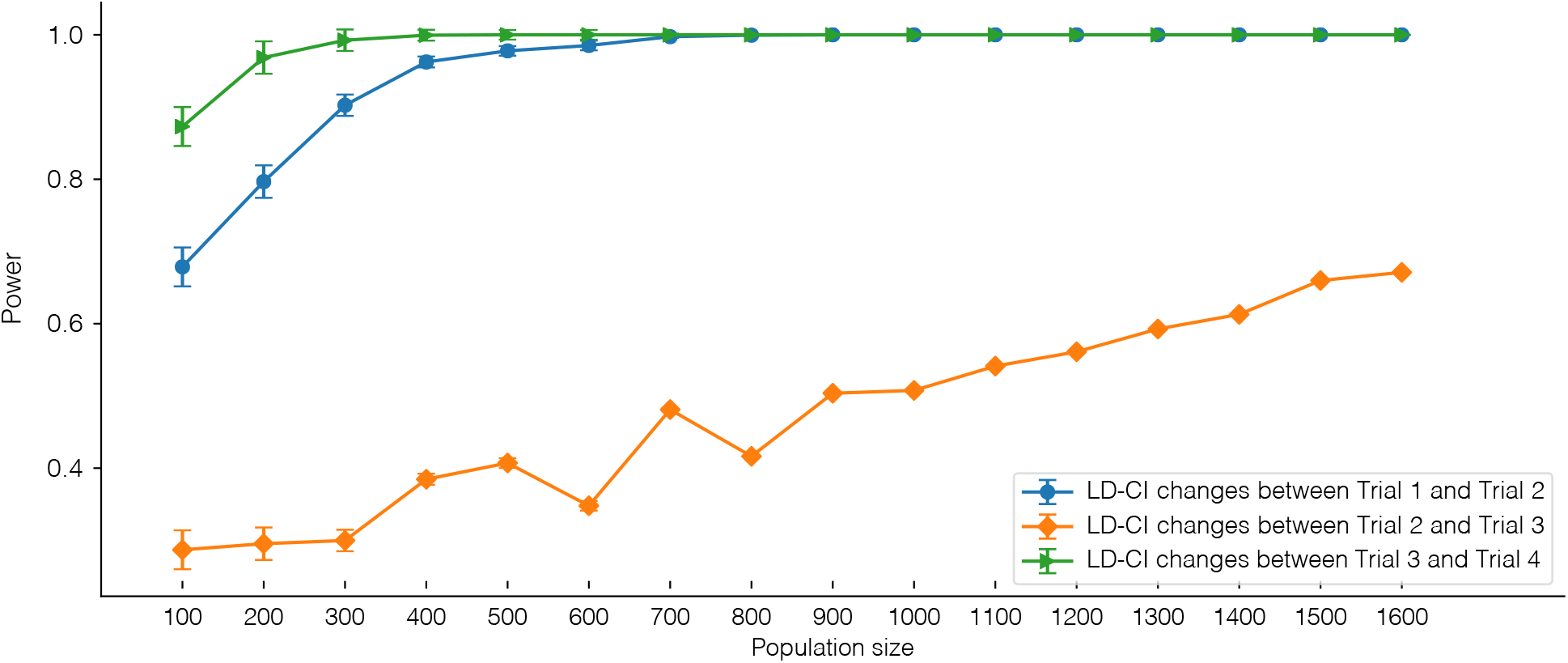
Power of detecting inter-trial behavioral differences improves with increased population size. Power for detecting the statistical significance of LD-CI changes over the trials was analyzed with population size ranging from 100 to 1600. Changes between Trial 1 and Trial 2 at 6dpf (blue) and between Trial 3 and Trial 4 at 7dpf (green) became exponentially detectable with the increment of the population size and plateaued after the size reached 500. For the detection of overnight decrease between Trial 3 and Trial 4 (orange), the power was linearly improved to less than 70% as the population size increased to the maximum.

## Discussion

Larval zebrafish have demonstrated learning and memory capabilities in both non-associative (Best et al., 2008; López-Schier, 2019; O’Neale et al., 2014; Randlett et al., 2019; Roberts et al., 2013; Wolman et al., 2011) and associative (Hinz et al., 2013; Lee et al., 2010; Valente et al., 2012) settings. In this study, using a large population of larval zebrafish, we have uncovered dynamic stimulus learning in two avoidance behaviors. The distinct patterns of changes in dark avoidance and center avoidance indicate that learning is stimulus specific. This observation also excludes other forces such as sensory adaptation or motor fatigue in driving behavioral changes, which should otherwise generalize across stimuli (Randlett et al., 2019; Rankin et al., 2009). The notion that learning drives avoidance behavioral changes is further supported by the observation that the locomotor activity (e.g., swim velocity) remains unchanged during habituation (**Figure S1**).

We have shown that short-term learning of dark avoidance starts with sensitization followed by habituation. While such a sequential occurrence of sensitization and habituation has been previously reported in rats (Davis, 1972; Geyer and Braff, 1987; Payne and Anderson, 1967; Szabo and Kolta, 1967) and in humans (Geyer and Braff, 1987; Ornitz and Guthrie, 1989), to our knowledge, this is the first demonstration of such a dynamic learning pattern in larval zebrafish. The fact that sensitization often occurs with an increased excitability of motor neurons (Thompson and Spencer, 1966) explains our observation of increased velocity in early time bins of each trial (**Figure S1**). In many cases, sensitization takes place during the first few presentations of a stimulus (McSweeney and Murphy, 2009). Our observation that sensitization precedes habituation in all four trials concords well with this notion. The initial predomination of sensitization over habituation is thought to represent the influence of a novel aversive stimulus on the central nervous system (Groves and Thompson, 1970; Ornitz and Guthrie, 1989) and is relevant to certain human disorders such as schizophrenia (Meincke et al., 2004).

Through iterative population subsampling, we have further shown that sensitization followed by habituation within a trial is detectable in a smaller cohort of 200 individuals. More importantly, the iterations enable us to quantify intra-trial learning performances with multiple indices. Remarkably, the duration of sensitization in each trial (STSD) mirrors the patterns of long-term habituation: a shortened STSD in the subsequent trial predicts inter-trial habituation whereas a lengthened STSD is linked to inter-trial (overnight) sensitization, a fascinating but not understood phenomenon that might involve sleep or active forgetting. Together, these findings suggest that the mechanisms that govern short-term (i.e., intratrial dynamics) and long-term (i.e., inter-trial dynamics) learning are interwoven.

Quantification of each larva’s long-term learning index (LTLI) has revealed individual differences in their patterns of learning. In general, the population displays a considerable spread distribution in long-term stimulus learning, which is also found in two recent studies (Pantoja et al., 2020; Randlett et al., 2019). However, our learning distribution shows both sensitization and habituation while only habituation is reported in these previous studies. In addition, the distribution curve deviates from a gaussian curve with an exceptionally high frequency at extreme learning indices for dark avoidance (**Figure 4B-D**), indicating a more divergent mechanism in shaping learning behaviors. A possible explanation for the relatively simple learning distribution in previous studies is that their learning performance is induced by intermittently delivered short-lived stimuli (Pantoja et al., 2020; Randlett et al., 2019). However, most stimuli in nature are stable over a longer period with more gradual transition of intensity. In addition to the mode of stimulus delivery, phenotyping paradigms used in previous studies usually restrain animals to a small well that partially immobilizes the animals, which potentially disrupts sensory feedback and alters neural responses (Stowers et al., 2017). In contrast, the light/dark preference paradigm used in this study allows observation of free-swimming larvae in the presence of stable stimuli, thereby providing a better mimicry of natural environment that helps uncovering stimulus learning patterns likely adopted for optimal survival in natural settings.

Contrary to a stereotyped learning dynamicity observed at the population level, an individual learns stimuli with more stochasticity. By constructing transition diagrams that describe the likelihood for an individual to display a subsequent learning type, we have uncovered a homeostatic control of learning dynamicity at the individual level (**Figure 4L-M**). Across successive trials, alternation between the two opposite learning types (sensitization and habituation) occurs with a probability about twice as high as that for the continuity of a same learning type. Homeostatic processes control biological parameters via negative feedback to restore the variable to a preadapted value, also termed as a set point (Knobil, 1999). Such a mechanism provides a plausible explanation for the alternating patterns between sensitization and habituation observed in our experiments.

It should also be noted that maintenance of a same learning type in successive trials has also been observed in small percentages of the population. These individuals might have a set point favored by an invariable learning style. Alternatively, this might be caused by a failure in homeostatic control, which is indicative of dysfunctionality in the underlying circuits. Further research focused on these “outlier” individuals could provide penetrating insights relevant to human disorders, which often represent extreme ends of a spectrum across the population.

In contrast to dark avoidance, center avoidance displays marginal changes within a trial (**Figure 2D**), low yet steady long-term habituation across trials without the phenomenon of overnight sensitization (**Figure 3B**). In addition, individual learning performance of center avoidance display a near normal distribution without significant deviations as is shown in the learning distribution of dark avoidance (**Figure 4B-D**). Despite these distinctions, individual long-term learning of center avoidance also displays alternating patterns of sensitization and habituation that is subjected to homeostatic control (**Figure 4M**). Taken together, although both dark avoidance and center avoidance are anxiety-related and controlled by a homeostatic process, distinct patterns of learning appear to be involved, the underlying mechanisms of which warrant further investigation.

Two previous studies employing the same light/dark preference paradigm fail to observe consistent behavioral changes within and across trials (Dahlén et al., 2019; Wagle et al., 2017). This is likely due to the fact that stimulus learning results in small but significant changes while displaying individual differences, making it difficult to detect with a small sample size. By subsampling the entire dataset, we have shown that a population of 200 is sufficient to detect short-term learning with initial sensitization followed by habituation. We have further demonstrated an exponential growth of the power for detecting long-term habituation when the sample size increases to 500. However, in order to uncover the overnight sensitization phenomenon, larger sample size is needed, as the power is less than 70% even with the maximized sample size of 1680 individuals. Thus, sufficient sample size is an important factor in studies of complex behaviors such as learning. Ongoing development in paradigm design and analytical tools will allow us to further increase throughput, which in turn should facilitate future mechanistic studies of genes and pathways involved in regulating the observed dynamic learning patterns.

In conclusion, our current study probes how larval zebrafish learn upon continuous exposure to a stable surrounding, mimicking possible encounters in nature. We uncover comprehensive patterns of stimulus learning, both short-term and long-term, and with stereotyped temporal dynamics and stimulus specificity at the population level. On an individual basis, learning exhibits stochasticity but is remarkably subjected to homeostatic control across stimuli. Some of these aspects require a sufficient sample size to unravel while others are tractable in smaller cohorts. These findings are well-suited for future investigation of underlying cellular and molecular mechanisms owing to the accessibility of larval zebrafish for *in vivo* functional imaging (Ahrens et al., 2013; Muto et al., 2013; Stewart et al., 2014) and high throughput molecular genetic studies (Chiu et al., 2016; Wolman et al., 2015)

## Methods

### Animals

Adult zebrafish (Danio rerio) used for larval production were bred in our facility at the University of California, San Francisco, CA and treated in accordance with IACUC regulations. After crossing, fertilized embryos were collected into petri dishes with blue egg water (0.12 g of CaSO4, 0.2 g of instant ocean salts from aquatic eco-systems, 30 ul of methylene blue in 1 L of H2O) and raised in a 28°C incubator for 2 days (**Figure 1A**). At the 3^rd^ day post fertilization (3dpf), dishes were transferred outside onto a light-blue colored surgical pad (VWR underpad Cat no. 82020-845) and exposed to a normal circadian cycle with 14h light and 10h dark period. On the day prior to behavior test (5dpf), larvae were individualized into twelve-well plates filled with 7ml egg water. All plates were kept under the same conditions.

### Behavior test

At 6dpf, dishes with individualized larvae were first moved to a same blue pad (luminance = 298 lux) in the behavior room one hour before test (pre-adaptation). Each larva was characterized for its light/dark preference and periphery/center preference (a.k.a thigmotaxis) with a previously established paradigm (Bai et al., 2016; Dahlén et al., 2019; Lau et al., 2011; Wagle et al., 2017). Briefly, an individual was transferred to a behavior chamber (4cm (L) × 4 cm (W) × 1.5 cm (H)) that was evenly divided into a light and a dark compartment by an opaque tape applied to the outside wall (**Figure 1A, top panel**). The border between the two sidewall compartments was aligned with clear and opaque black acrylic stripes placed underneath the transparent chamber; together, they were placed on a trans-illuminator so that light only pass through the clear (luminance = 576 lux) but not the opaque black acrylic stripes (luminance = 0 lux). For behavior recording, a camera (Panasonic) with infrared filters (ACRYLITE IR acrylic 11460) was fixed over the chamber and was connected to the software Noldus MPEG recorder 2.1. A group of sixteen chambers were recorded at a time for an 8-min trial and up to 192 individuals were tested in a 2hr session (**Figure 1A, middle panel**). An early trial was performed from 9AM to 11AM followed by a later trial from 11AM to 1PM at 6dpf, leaving a 2hr interval between the two trials of a same individual. Same procedure was repeated at 7dpf for a total of four trials. This experiment assay was weekly conducted to test a total of 1680 larvae. Behavior recordings were digitized by EthoVision XT 13 to produce raw data output for further analysis.

## Data analysis

All data analysis was performed with custom code written in Python (Supplemental code). The Light/Dark Choice Index (LD-CI) and Periphery-Center Choice index (PC-CI) was calculated using the equations described in Dahlén et al.(2019) (**Figure 1B**). Increase of each index indicates an increased time an animal spent in the area to which it was initially aversive (i.e. dark zone for LD-CI and inner zone for PC-CI). These two indices were calculated for each trial as well as for sixteen non-overlapped 30s-periods within a trial, producing a trial index and a bin index for each individual (Supplemental **Table S1 and S2**).

Population mean of bin indices was calculated and plotted chronologically to illustrate within-trial kinetics (**Figure 2A and 2D**). The resulted graph indicated two sequentially occurred periods within each trial: sensitization and habituation. To characterize these two periods, we randomly subsampled 200 individuals and found the bin with lowest mean LD-CI as the one that divides the period of sensitization from that of habituation. We defined the period (s) from the beginning to the middle point of the divider bin as the short-term sensitization duration (STSD) and the remaining time of the trial as the habituation duration. Using an equation (Equation 1.1) derived from Raymond et al. (2012), a short-term sensitization index (STSI) was calculated for each subsampled data consisting of 200 individuals to indicate a percentage reduction of bin LD-CI over the sensitized period. Similarly, a short-term habituation index (STHI) was computed by comparing percentage change of bin LD-CI across the habituated period (Equation 1.2). By iterating the subsampling process for 1000 runs, we generated vectors of STSD (s), STSI (%) and STHI (%) of each trial followed by inter-trial comparisons using paired one-tailed t-test (**Figure 2B-C**).

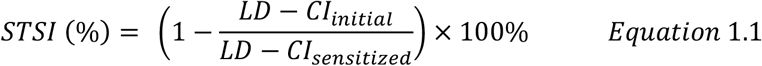

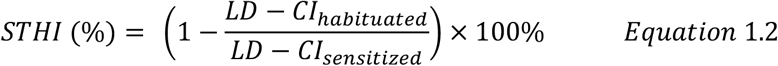

The means of trial indices across all individuals were compared with a paired one-tailed t-test among the four trials to reveal inter-trial changes (**Figure 3A-B**). For every pair of two successive trials, a significantly higher index of the subsequent trial compared to that of the previous trial indicates a habituation to the aversiveness of the stimulus at the population level. On the other hand, a later trial with a lower index suggests a sensitization to the stimulus. Three other LD-CI related components including number of dark zone entries, average duration of dark zone entry, and the latency to the first dark zone entry, were also calculated for inter-trial comparisons (**Figure 3C-E**).

The long-term learning index (LTLI) was calculated for individual larva to indicate a change of aversiveness in the later trial relative to early trial using Equation 2. To make the distribution comparable across the different behavior components, a min-max normalization (Juszczak et al., 2002) was used before plugging values into the equation. When calculating latency, value of early trial instead of later trial is used as the numerator in Equation 2 so that habituation of all the behavior components is always indicated by a value greater than 1. By contrast, a LTLI lower than 1 suggests an individual sensitized to the stimuli. Distribution of LTLI for each behavioral component is presented using a kernel density curve implemented by Scott’s Rule (Scott, 2015) (**Figure 4B-D**).

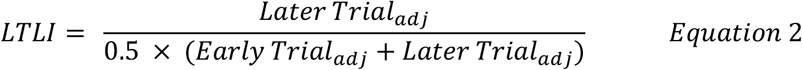

As is indicated by Equation 2, LTLI ranges from 0 to 2 and the two limits are reached when a minimum value −1 is detected in one of two successive trials, resulting in either Later Trial_a_dj or Early Trial_a_dj being 0. Individuals in such a scenario would have a fixed LTLI of either 0 or 2 regardless of their true learning performance. Therefore, these LTLIs cannot properly reflect the learning behavior and excluded for plotting the distribution curve. However, these individuals are still categorized into the group of habituation or sensitization based on calculating their inter-trial differences. Moreover, individuals with a minimum value −1 in both trials are categorized as non-learners (LTHI = 1). Portion of each group is presented as a pie chart for learning of dark avoidance (**Figure 4E-G**) and center avoidance (**Figure 4H-J**).

Based on the differences of LD-CI in two successive trials, each individual was assigned a certain learning type, resulting in three learning types for each individual across four trials (**Figure 4K**). We then calculated the probability of learning type switching to analyze learning dynamicity. For larvae with a given learning type (e.g., S), we computed the portion with the same learning type (e.g., S) after a transition to derive the probability of maintaining learning consistency. We also computed the portion with a different learning type (e.g., H) after a transition to indicate the probability of learning type switching. The resulting two sets of probabilities were summarized in a transition diagram to illustrate learning dynamicity at each transition: solid arrows indicate majority while dashed arrows indicate minority; numbers in black indicated transition 1 while numbers in blue denoted transition 2 (**Figure 4L-M**).

For the impact of population size on the detection of long-term habituation, we subsampled data of different sizes ranging from 100 to 1600. For a given size n, we randomly selected a subset of n individuals from the entire data and performed paired one-tailed t-test between the trials to compute a power value using ‘power_ttest’ function in a python package ‘pingouin’ (Vallat, 2018). The process was iterated for 100 times to produce a mean value as the ultimate estimate of the power corresponding to the sample size n. In this way, we recorded power changes across 16 different sample sizes (**Figure 5**).

## Acknowledgements

We thank the Guo lab members in particular Mahendra Wagle, for technical advice, suggestions, and helpful discussions, and Michael Munchua and Vivian Yuan for excellent fish care. This work was supported by NIH R01GM132500 (to S.G.), and associated diversity supplement award (to R.C.).

## Author contributions

Jiale Xu, Conceptualization, Animal curation, Data management, Formal analysis, Investigation; Romelo Casanave, Animal curation, Behavior recording; Guo Su, Conceptualization, Formal analysis, Supervision

## Competing interests

The authors declare that they have no competing interests.

## Supplemental Figures

**Figure S1.**
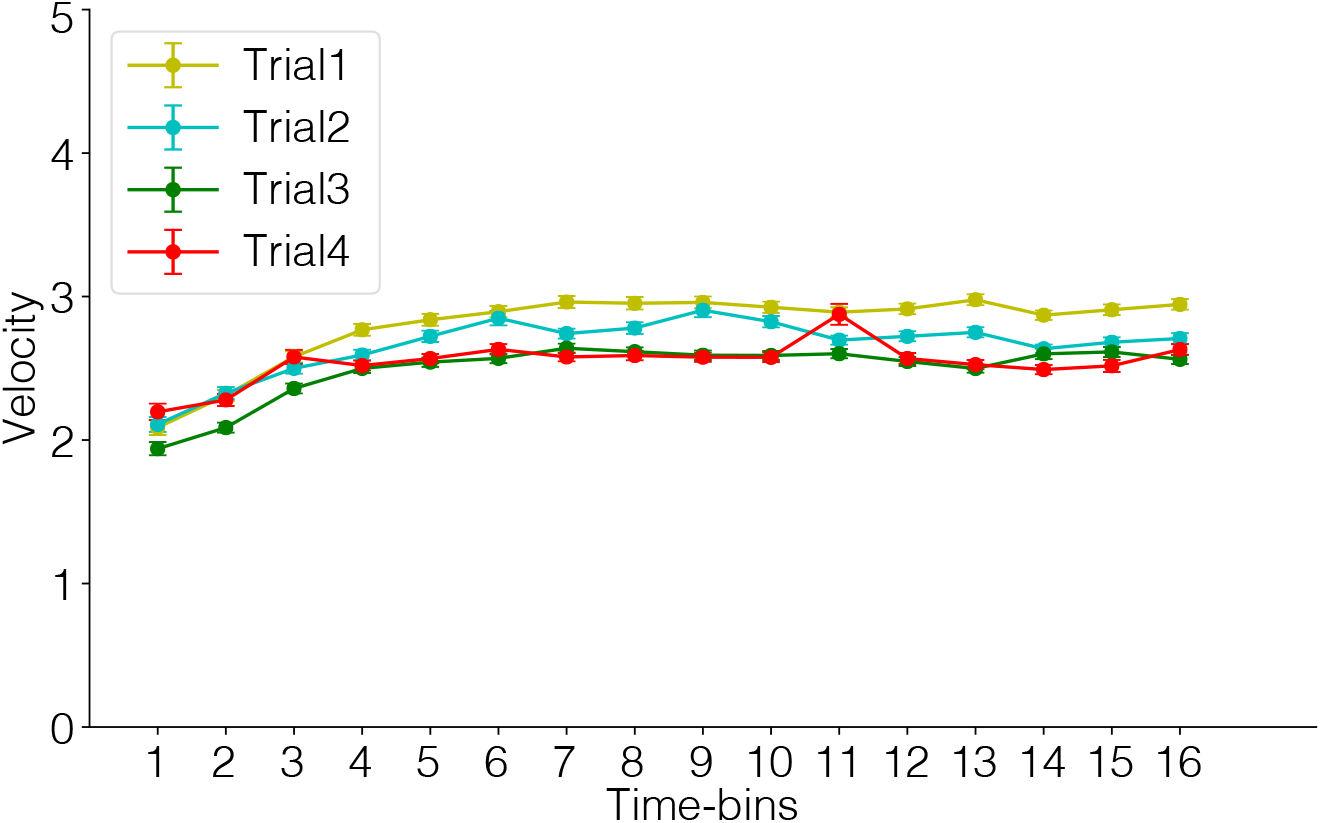
Intra-trial dynamics of velocity. Within each trial, mean velocity across all the larvae (n=1680) was increased during the first four bins, which correspond to the sensitization period. After that, mean velocity plateaued with slight fluctuations throughout the rest of the trial when individuals underwent habituation.

**Figure S2.**
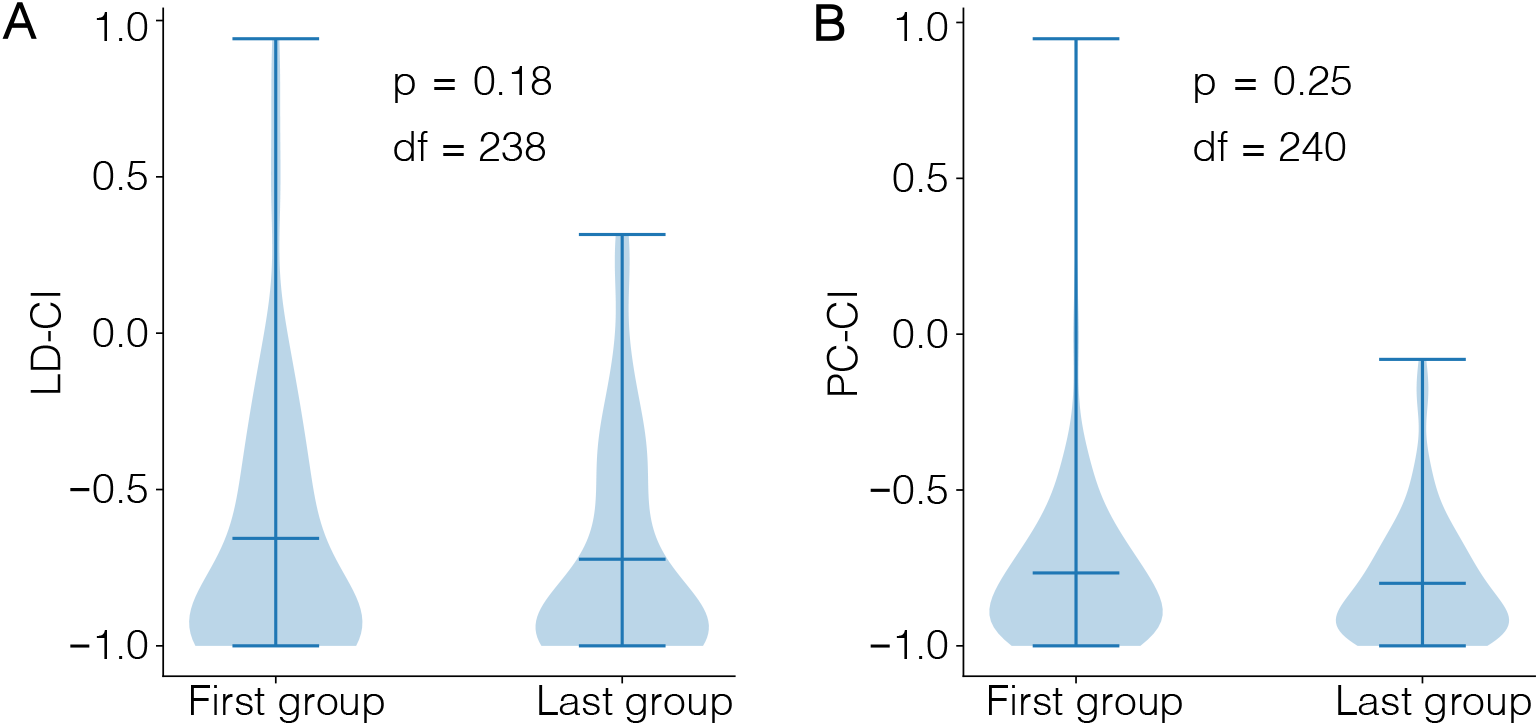
Comparisons of Trial 1 means between the first and last tested groups. Comparing Trial 1 means between the first and last tested groups (unpaired twotailed t-test, n_first_=143, n_last_=101), tested with ~ 2hrs apart, resulted insignificant differences, confirming that the observed long-term habituation effect in dark avoidance (A) and center avoidance (B) is not due to experimental time.

## Notes

### Competing Interest Statement

The authors have declared no competing interest.

