## Supplemental Code for "Larval zebrafish display dynamic learning of aversive stimuli in a constant visual surrounding"


In [1]:

```
import pandas as pd
import matplotlib as mpl
import matplotlib.pyplot as plt
import numpy as np
import pingouin as pg
import scipy.stats as stats
from matplotlib.lines import Line2D
import matplotlib.font_manager as font_manager
from matplotlib import rc
import math
```

In [7]:

```
mpl.matplotlib_fname()
```

Out[7]:

```
'/Users/jialexu/opt/anaconda3/lib/python3.7/site-packages/matplotlib/mpl-data/matplotlibrc'
```

In [10]:

```
##Intra-trial LD-CI dynamics##

#data import#
df_accumulative_pb_ci=pd.read_excel('Supplemental_Table_S1.xlsx', 
                      sheet_name='Binned_LD-CI', index_col=0, header=3)

#data clean#
df_t1_ci=df_accumulative_pb_ci.dropna(axis=0).loc[:,df_accumulative_pb_ci.columns.str.contains('1-')].transpose()
df_t2_ci=df_accumulative_pb_ci.dropna(axis=0).loc[:,df_accumulative_pb_ci.columns.str.contains('2-')].transpose()
df_t3_ci=df_accumulative_pb_ci.dropna(axis=0).loc[:,df_accumulative_pb_ci.columns.str.contains('3-')].transpose()
df_t4_ci=df_accumulative_pb_ci.dropna(axis=0).loc[:,df_accumulative_pb_ci.columns.str.contains('4-')].transpose()

#data visulization#
font ={'family' : 'Arial',
      'size' : 18}
mpl.rc('font', **font)
fig, ax = plt.subplots(figsize=(7,7), sharey=True)
df_t1_ci.mean(axis=1).plot(marker='o',yerr=df_t1_ci.sem(axis=1), capsize=4, label='Trial1', color='y', ax=ax)
df_t2_ci.mean(axis=1).plot(marker='o',yerr=df_t2_ci.sem(axis=1), capsize=4, label='Trial2', color='c', ax=ax)
df_t3_ci.mean(axis=1).plot(marker='o',yerr=df_t3_ci.sem(axis=1), capsize=4, label='Trial3', color='g', ax=ax)
df_t4_ci.mean(axis=1).plot(marker='o',yerr=df_t4_ci.sem(axis=1), capsize=4, label='Trial4', color='r', ax=ax)
ax.spines['top'].set_visible(False)
ax.spines['right'].set_visible(False)
ax.set_xticks(np.arange(0,16))
ax.set_xticklabels(labels=np.arange(1,17))
ax.set_xlim(-1,17)
ax.set_ylim(-1,-0.3)

ax.set_ylabel('LD-CI')
ax.set_xlabel('Time-bins')
ax.legend(loc='upper left')


#plt.show()
plt.savefig('fig2A.pdf',bbox_inches = 'tight',
    pad_inches = 0)
```

In [13]:

```
df_accumulative_pb_velocity=pd.read_excel('Supplemental_Table_S1.xlsx', 
                      sheet_name='Binned_Velocity', index_col=0, header=3)

df_t1_velocity=df_accumulative_pb_velocity.dropna(axis=0).loc[:,df_accumulative_pb_velocity.columns.str.contains('1-')].transpose()
df_t2_velocity=df_accumulative_pb_velocity.dropna(axis=0).loc[:,df_accumulative_pb_velocity.columns.str.contains('2-')].transpose()
df_t3_velocity=df_accumulative_pb_velocity.dropna(axis=0).loc[:,df_accumulative_pb_velocity.columns.str.contains('3-')].transpose()
df_t4_velocity=df_accumulative_pb_velocity.dropna(axis=0).loc[:,df_accumulative_pb_velocity.columns.str.contains('4-')].transpose()
#drop individuals with velocity equal to '-'
df_t1_velocity=df_t1_velocity.drop(columns=df_t1_velocity.columns[(df_t1_velocity=='-').any()])
df_t2_velocity=df_t2_velocity.drop(columns=df_t2_velocity.columns[(df_t2_velocity=='-').any()])
df_t3_velocity=df_t3_velocity.drop(columns=df_t3_velocity.columns[(df_t3_velocity=='-').any()])
df_t4_velocity=df_t4_velocity.drop(columns=df_t4_velocity.columns[(df_t4_velocity=='-').any()])

#data visualization#
font ={'family' : 'Arial',
      'size' : 20}
mpl.rc('font', **font)
fig, ax = plt.subplots(figsize=(10,6), sharey=True)
df_t1_velocity.mean(axis=1).plot(marker='o',yerr=df_t1_velocity.sem(axis=1), capsize=4, label='Trial1', color='y', ax=ax)
df_t2_velocity.mean(axis=1).plot(marker='o',yerr=df_t2_velocity.sem(axis=1), capsize=4, label='Trial2', color='c', ax=ax)
df_t3_velocity.mean(axis=1).plot(marker='o',yerr=df_t3_velocity.sem(axis=1), capsize=4, label='Trial3', color='g', ax=ax)
df_t4_velocity.mean(axis=1).plot(marker='o',yerr=df_t4_velocity.sem(axis=1), capsize=4, label='Trial4', color='r', ax=ax)
ax.spines['top'].set_visible(False)
ax.spines['right'].set_visible(False)
ax.set_xticks(np.arange(0,16))
ax.set_xticklabels(labels=np.arange(1,17))
ax.set_yticklabels(labels=np.arange(0,6))
ax.set_xlim(-1,17)
ax.set_ylim(0,5)
ax.set_ylabel('Velocity')
ax.set_xlabel('Time-bins')
ax.legend(loc='upper left')
plt.savefig('figS1.pdf',bbox_inches = 'tight',
    pad_inches = 0)
plt.show()
```

In [15]:

```
##Subsample iteration with cohorts of 200 individuals##

import random
font ={'family' : 'Arial'}
mpl.rc('font', **font)
fig, ax=plt.subplots(1,4,figsize=(12,9), sharey=True,)
color=['y', 'c', 'g', 'r']
for i, df in enumerate ([df_t1_ci,df_t2_ci,df_t3_ci,df_t4_ci]):  
    meandic={}
    for j in range (0, 1000):
            subset=random.sample(range(0,1680), 200)
            meandic.update({j:df.iloc[:,subset].mean(axis=1)})
    df_mean=pd.DataFrame(meandic)
    df_summary=pd.DataFrame({'mean':df_mean.mean(axis=1), 'std':df_mean.std(axis=1)})
    ax[i].fill_between(np.arange(0,16), 
                       df_summary['mean']+1.96*df_summary['std'], df_summary['mean']-1.96*df_summary['std'],
                      color=color[i], label='95%CI')
    
    ax[i].plot(df_summary['mean'], color='b', label='mean')
    ax[i].legend(loc='upper left',fontsize=20) 
    ax[i].set_xticklabels(labels=np.arange(1,17), rotation=30,fontsize=11)
    ax[i].set_ylim(-1, -0.1)
    ax[i].spines['right'].set_visible(False)
    ax[i].spines['top'].set_visible(False) 
    ax[i].set_xlabel('Trial'+' '+str(i+1),fontsize=20)
    ax[i].set_ylabel('LD-CI',fontsize=20)
    plt.rcParams['ytick.labelsize']=20

plt.savefig('fig2B.pdf',bbox_inches = 'tight',
    pad_inches = 0)
```

In [19]:

```
## Statistical analysis of the resulted 1000 cohorts of 200 individuals##


def intratrial(df, subset):

    df_new=df.iloc[:, subset]
    key=df_new.mean(axis=1).sort_values().index[0]
    sensdur=int(key.split('-')[1][3])*60+int(key.split('-')[1][5:7])+15   
    sensitized_ci=df_new.loc[key].mean()
    initial_ci=df_new.iloc[0].mean()
    habituated_ci=df_new.iloc[15].mean()
        
    STSI=(1-initial_ci/sensitized_ci)
    STSI=STSI*100
    STHI=(1-habituated_ci/sensitized_ci)
    STHI=STHI*100    
    return pd.Series([sensdur, STSI, STHI])

sensdur_list=[]
STSI_list=[]
STHI_list=[]
import random
for i in range (0,1000):
    subset=random.sample(range(0,1680), 200)
    t1_sensdur=intratrial(df_t1_ci, subset)[0]
    t2_sensdur=intratrial(df_t2_ci, subset)[0]
    t3_sensdur=intratrial(df_t3_ci, subset)[0]
    t4_sensdur=intratrial(df_t4_ci, subset)[0]
    sensdur=pd.Series([t1_sensdur,t2_sensdur,t3_sensdur,t4_sensdur], name=i)
    sensdur_list.append(sensdur)
    
    t1_STSI=intratrial(df_t1_ci, subset)[1]
    t2_STSI=intratrial(df_t2_ci, subset)[1]
    t3_STSI=intratrial(df_t3_ci, subset)[1]
    t4_STSI=intratrial(df_t4_ci, subset)[1]
    STSI=pd.Series([t1_STSI,t2_STSI,t3_STSI,t4_STSI], name=i)
    STSI_list.append(STSI)

    t1_STHI=intratrial(df_t1_ci, subset)[2]
    t2_STHI=intratrial(df_t2_ci, subset)[2]
    t3_STHI=intratrial(df_t3_ci, subset)[2]
    t4_STHI=intratrial(df_t4_ci, subset)[2]
    STHI=pd.Series([t1_STHI,t2_STHI,t3_STHI,t4_STHI], name=i)
    STHI_list.append(STHI)
    

df_sensdur=pd.concat(sensdur_list,axis=1)
df_sensdur.index=['Trial1', 'Trial2', 'Trial3', 'Trial4']
df_sensdur=df_sensdur.transpose()
df_STSI=pd.concat(STSI_list,axis=1)
df_STSI.index=['Trial1', 'Trial2', 'Trial3', 'Trial4']
df_STSI=df_STSI.transpose()
df_STHI=pd.concat(STHI_list,axis=1)
df_STHI.index=['Trial1', 'Trial2', 'Trial3', 'Trial4']
df_STHI=df_STHI.transpose()

fig,ax=plt.subplots(3,1, figsize=(7,10))
ylabels=['STSD (sec)', 'STSI (%)', 'STHI (%)']
yticks=[np.arange(0,120, 20), np.arange(0,12,2), np.arange(0,70,10)]
plt.subplots_adjust(hspace=0.5)
for i, df in enumerate ([df_sensdur,df_STSI,df_STHI]):
    ax[i].bar(np.array([1,2,4,5])+0.35, df.mean(), 0.35, yerr=df.sem(),capsize=4, color=['y', 'c', 'g', 'r'])
    ax[i].set_xticks(np.array([1,2,4,5]) + 0.35)
    ax[i].set_xticklabels(('Trial1', 'Trial2', 'Trial3', 'Trial4') ,fontsize=20)
    ax[i].set_yticks(yticks[i])
    ax[i].spines['right'].set_visible(False)
    ax[i].spines['top'].set_visible(False)
    ax[i].set_ylabel(ylabels[i],fontsize=20)
    ax[i].set_ylim(0, df.mean().max()*1.2)
    result1=pg.ttest(df['Trial1'], df['Trial2'],paired=True, tail='one-sided').round(2)
    result2=pg.ttest(df['Trial2'], df['Trial3'],paired=True, tail='one-sided').round(2)
    result3=pg.ttest(df['Trial3'], df['Trial4'],paired=True, tail='one-sided').round(2)
    result=pd.concat([result1,result2,result3])
    pvaluelist=result.iloc[:,3]
    symbollist=[]
    for p in pvaluelist:
        if p>0 and p<0.5:
            symbol='*'
        elif p<0.01:
            symbol='**'
        else:
            symbol='ns'
        symbollist.append(symbol)
    difft2t1=round((df['Trial2']-df['Trial1']).mean(),2)
    difft3t2=round((df['Trial3']-df['Trial2']).mean(),2)
    difft4t3=round((df['Trial4']-df['Trial3']).mean(),2)
    ax[i].annotate(str(difft2t1)+symbollist[0], xy=(0.17, 0.95), xytext=(0.17, 1), xycoords='axes fraction', 
                ha='center', va='bottom',
                arrowprops=dict(arrowstyle='-[, widthB=2, lengthB=0.2', lw=1.0), fontsize=20)
    ax[i].annotate(str(difft3t2)+symbollist[1], xy=(0.5, 0.95), xytext=(0.5, 1), xycoords='axes fraction', 
                ha='center', va='bottom',
                arrowprops=dict(arrowstyle='-[, widthB=4.0, lengthB=0.2', lw=1.0),fontsize=20)
    ax[i].annotate(str(difft4t3)+symbollist[2], xy=(0.82, 0.95), xytext=(0.82, 1), xycoords='axes fraction', 
                ha='center', va='bottom',
                arrowprops=dict(arrowstyle='-[, widthB=2, lengthB=0.2', lw=1.0),fontsize=20)

plt.savefig('fig2C.pdf',bbox_inches = 'tight',
    pad_inches = 0)
```

In [21]:

```
##Intra-trial LD-CI dynamics##

#data import#
df_accumulative_pb_th=pd.read_excel('Supplemental_Table_S1.xlsx', 
                      sheet_name='Binned_PC-CI', index_col=0, header=3)

fig, ax = plt.subplots(figsize=(8,8), sharey=True)

#data clean#
df_t1_th=df_accumulative_pb_th.dropna(axis=0).loc[:,df_accumulative_pb_th.columns.str.contains('1-')].transpose()
df_t2_th=df_accumulative_pb_th.dropna(axis=0).loc[:,df_accumulative_pb_th.columns.str.contains('2-')].transpose()
df_t3_th=df_accumulative_pb_th.dropna(axis=0).loc[:,df_accumulative_pb_th.columns.str.contains('3-')].transpose()
df_t4_th=df_accumulative_pb_th.dropna(axis=0).loc[:,df_accumulative_pb_th.columns.str.contains('4-')].transpose()

#data visualization#
df_t1_th.mean(axis=1).plot(marker='o',yerr=df_t1_th.sem(axis=1), capsize=4, label='Trial1',color='y', ax=ax)
df_t2_th.mean(axis=1).plot(marker='o',yerr=df_t2_th.sem(axis=1), capsize=4, label='Trial2',color='c', ax=ax)
df_t3_th.mean(axis=1).plot(marker='o',yerr=df_t3_th.sem(axis=1), capsize=4, label='Trial3',color='g', ax=ax)
df_t4_th.mean(axis=1).plot(marker='o',yerr=df_t4_th.sem(axis=1), capsize=4, label='Trial4',color='r', ax=ax)
ax.set_xticks(np.arange(0,16))
ax.set_xticklabels(labels=np.arange(1,17),fontsize=15)
ax.set_xlim(-1,17)
ax.spines['top'].set_visible(False)
ax.spines['right'].set_visible(False)
ax.set_xlabel('Time-bins', fontsize=20)
ax.set_ylabel('PC-CI', fontsize=20)
ax.set_xlim(-1,17)
ax.set_ylim(-1,-0.3)
ax.legend(loc='upper left',fontsize=20)

plt.savefig('fig2D.pdf',bbox_inches = 'tight',
    pad_inches = 0)
plt.show()
```

In [22]:

```
##Inter-trial comparions of LD-CI##

#data import#
df_pop_arena_trial=pd.read_excel('Supplemental_Table_S2.xlsx', 
                    index_col=0, header=1)

#data analysis#
df=df_pop_arena_trial
result1=pg.ttest(df.loc[df['Trial']==2].iloc[:,3], 
                df.loc[df['Trial']==1].iloc[:,3], 
                paired=True, tail='one-sided').round(2)
result2=pg.ttest(df.loc[df['Trial']==3].iloc[:,3], 
                df.loc[df['Trial']==2].iloc[:,3], 
                paired=True, tail='one-sided').round(2)
result3=pg.ttest(df.loc[df['Trial']==4].iloc[:,3], 
                df.loc[df['Trial']==3].iloc[:,3], 
                paired=True, tail='one-sided').round(2)
result=pd.concat([result1,result2,result3])

difft2t1=round(df.loc[df['Trial']==2].iloc[:,3].mean()-df.loc[df['Trial']==1].iloc[:,3].mean(),2)
difft3t2=round(df.loc[df['Trial']==3].iloc[:,3].mean()-df.loc[df['Trial']==2].iloc[:,3].mean(),2)
difft4t3=round(df.loc[df['Trial']==4].iloc[:,3].mean()-df.loc[df['Trial']==3].iloc[:,3].mean(),2)

pvaluelist=result.iloc[:,3]
symbollist=[]
for p in pvaluelist:
    if p>0 and p<0.5:
        symbol='*'
    elif p<0.01:
        symbol='**'
    else:
        symbol='ns'
    symbollist.append(symbol)

#data visualize#
fig, ax = plt.subplots(figsize=(8,6))
parts=ax.violinplot([df.loc[df['Trial']==1].iloc[:,3],
               df.loc[df['Trial']==2].iloc[:,3],
               df.loc[df['Trial']==3].iloc[:,3],
               df.loc[df['Trial']==4].iloc[:,3]], showmeans=True, positions=[1,2,4,5])
ax.spines['right'].set_visible(False)
ax.spines['top'].set_visible(False)

plt.xticks(np.r_[1,2,4,5], labels=['Trial 1', 'Trial 2','Trial 3','Trial 4'], fontsize=18)
colors=['y', 'c', 'g', 'r']
for pc,color in zip(parts['bodies'], colors):
        pc.set_color(color)
plt.ylim(-1.1,1.5)


plt.annotate(str(difft2t1)+symbollist[0], xy=(0.2, 0.85), xytext=(0.2, 0.9), xycoords='axes fraction', 
            ha='center', va='bottom',fontsize=18,
            arrowprops=dict(arrowstyle='-[, widthB=2.0, lengthB=0.2', lw=1.0))
plt.annotate(str(difft3t2)+symbollist[1], xy=(0.5, 0.85), xytext=(0.5, 0.9), xycoords='axes fraction', 
            ha='center', va='bottom',fontsize=18,
            arrowprops=dict(arrowstyle='-[, widthB=5.0, lengthB=0.2', lw=1.0))
plt.annotate(str(difft4t3)+symbollist[2], xy=(0.8, 0.85), xytext=(0.8, 0.9), xycoords='axes fraction', 
            ha='center', va='bottom',fontsize=18,
            arrowprops=dict(arrowstyle='-[, widthB=2.0, lengthB=0.2', lw=1.0))

plt.annotate('2h interval', xy=(0.2, 0.4), xytext=(0.2, 0.4), xycoords='axes fraction', 
            ha='center', va='bottom',fontsize=18)
plt.annotate('6dpf', xy=(0.2, 0.5), xytext=(0.2, 0.5), xycoords='axes fraction', 
            ha='center', va='bottom',fontsize=18)
plt.annotate('2h interval', xy=(0.8, 0.4), xytext=(0.8, 0.4), xycoords='axes fraction', 
            ha='center', va='bottom',fontsize=18)
plt.annotate('7dpf', xy=(0.8, 0.5), xytext=(0.8, 0.5), xycoords='axes fraction', 
            ha='center', va='bottom',fontsize=18)
plt.annotate('22h interval', xy=(0.5, 0.4), xytext=(0.5, 0.4), xycoords='axes fraction', 
            ha='center', va='bottom',fontsize=18)
plt.annotate('6dpf~7dpf', xy=(0.5, 0.5), xytext=(0.5, 0.5), xycoords='axes fraction', 
            ha='center', va='bottom',fontsize=18)


plt.ylabel('LD-CI', fontsize=18)
plt.savefig('fig3A.pdf',bbox_inches = 'tight',
    pad_inches = 0)
plt.show()
```

In [23]:

```
#data analysis#
df=df_pop_arena_trial
result1=pg.ttest(df.loc[df['Trial']==2].iloc[:,4], 
                df.loc[df['Trial']==1].iloc[:,4], 
                paired=True, tail='one-sided').round(2)
result2=pg.ttest(df.loc[df['Trial']==3].iloc[:,4], 
                df.loc[df['Trial']==2].iloc[:,4], 
                paired=True, tail='one-sided').round(2)
result3=pg.ttest(df.loc[df['Trial']==4].iloc[:,4], 
                df.loc[df['Trial']==3].iloc[:,4], 
                paired=True, tail='one-sided').round(2)
result=pd.concat([result1,result2,result3])

difft2t1=round(df.loc[df['Trial']==2].iloc[:,4].mean()-df.loc[df['Trial']==1].iloc[:,4].mean(),2)
difft3t2=round(df.loc[df['Trial']==3].iloc[:,4].mean()-df.loc[df['Trial']==2].iloc[:,4].mean(),2)
difft4t3=round(df.loc[df['Trial']==4].iloc[:,4].mean()-df.loc[df['Trial']==3].iloc[:,4].mean(),2)

pvaluelist=result.iloc[:,3]
symbollist=[]
for p in pvaluelist:
    if p>0.01 and p<0.05:
        symbol='*'
    elif p<0.01:
        symbol='**'
    else:
        symbol='ns'
    symbollist.append(symbol)
    
#data visualize#
fig, ax = plt.subplots(figsize=(8,6))
parts=ax.violinplot([df.loc[df['Trial']==1].iloc[:,4],
               df.loc[df['Trial']==2].iloc[:,4],
               df.loc[df['Trial']==3].iloc[:,4],
               df.loc[df['Trial']==4].iloc[:,4]], showmeans=True, positions=[1,2,4,5])
ax.spines['right'].set_visible(False)
ax.spines['top'].set_visible(False)

plt.xticks(np.r_[1,2,4,5], labels=['Trial 1', 'Trial 2','Trial 3','Trial 4'], fontsize=18)
colors=['y', 'c', 'g', 'r']
for pc,color in zip(parts['bodies'], colors):
        pc.set_color(color)
plt.ylim(-1.1,1.5)


plt.annotate(str(difft2t1)+symbollist[0], xy=(0.2, 0.85), xytext=(0.2, 0.9), xycoords='axes fraction', 
            ha='center', va='bottom',fontsize=18,
            arrowprops=dict(arrowstyle='-[, widthB=2.5, lengthB=0.2', lw=1.0))
plt.annotate(str(difft3t2)+symbollist[1], xy=(0.5, 0.85), xytext=(0.5, 0.9), xycoords='axes fraction', 
            ha='center', va='bottom',fontsize=18,
            arrowprops=dict(arrowstyle='-[, widthB=5.0, lengthB=0.2', lw=1.0))
plt.annotate(str(difft4t3)+symbollist[2], xy=(0.8, 0.85), xytext=(0.8, 0.9), xycoords='axes fraction', 
            ha='center', va='bottom',fontsize=18,
            arrowprops=dict(arrowstyle='-[, widthB=2.5, lengthB=0.2', lw=1.0))

plt.annotate('2h interval', xy=(0.2, 0.4), xytext=(0.2, 0.4), xycoords='axes fraction', 
            ha='center', va='bottom',fontsize=18)
plt.annotate('6dpf', xy=(0.2, 0.5), xytext=(0.2, 0.5), xycoords='axes fraction', 
            ha='center', va='bottom',fontsize=18)
plt.annotate('2h interval', xy=(0.8, 0.4), xytext=(0.8, 0.4), xycoords='axes fraction', 
            ha='center', va='bottom',fontsize=18)
plt.annotate('7dpf', xy=(0.8, 0.5), xytext=(0.8, 0.5), xycoords='axes fraction', 
            ha='center', va='bottom',fontsize=18)
plt.annotate('22h interval', xy=(0.5, 0.4), xytext=(0.5, 0.4), xycoords='axes fraction', 
            ha='center', va='bottom',fontsize=18)
plt.annotate('6dpf~7dpf', xy=(0.5, 0.5), xytext=(0.5, 0.5), xycoords='axes fraction', 
            ha='center', va='bottom',fontsize=18)


plt.ylabel('PC-CI', fontsize=18)
plt.savefig('fig3B.pdf',bbox_inches = 'tight',
    pad_inches = 0)
plt.show()
```

In [24]:

```
#data analysis#

df=df_pop_arena_trial
fig, ax = plt.subplots(1,3, figsize=(23,7))
ylabels=['Number of dark zone entry', 'Average duration of dark zone entry', 'Latency to 1st dark zone entry']

#introduce a function to exclude outliers
def exoutliers(df, parameter):
    Q1=df[parameter].quantile(0.25)
    Q3=df[parameter].quantile(0.75)
    IQR=Q3-Q1
    return df.loc[(df[parameter]>(Q1-1.75*IQR))&(df[parameter]<(Q3+1.75*IQR))]

for i, parameter in enumerate (ylabels):
    result1=pg.ttest(df.loc[df['Trial']==2][parameter], 
                df.loc[df['Trial']==1][parameter], 
                paired=True, tail='one-sided').round(2)
    result2=pg.ttest(df.loc[df['Trial']==3][parameter], 
                df.loc[df['Trial']==2][parameter], 
                paired=True, tail='one-sided').round(2)
    result3=pg.ttest(df.loc[df['Trial']==4][parameter], 
                df.loc[df['Trial']==3][parameter], 
                paired=True, tail='one-sided').round(2)
    result=pd.concat([result1,result2,result3])
    
#data visualize#

    df_t1=exoutliers(df.loc[df['Trial']==1], parameter)
    df_t2=exoutliers(df.loc[df['Trial']==2], parameter)
    df_t3=exoutliers(df.loc[df['Trial']==3], parameter)
    df_t4=exoutliers(df.loc[df['Trial']==4], parameter)
    
    parts=ax[i].violinplot([df_t1[parameter],df_t2[parameter],
                            df_t3[parameter],df_t4[parameter]], showmeans=True, positions=[1,2,4,5])
    
    
    
    ax[i].spines['right'].set_visible(False)
    ax[i].spines['top'].set_visible(False)
    ax[i].set_xticks([1,2,4,5])
    ax[i].set_xticklabels(labels=['Trial 1', 'Trial 2','Trial 3','Trial 4'],fontsize=20)
    colors=['y', 'c', 'g', 'r']
    for pc,color in zip(parts['bodies'], colors):
            pc.set_color(color)
    ax[i].set_ylim(df_t4[parameter].max()*(-0.02),df_t4[parameter].max()*1.2)

    difft2t1=round(df.loc[df['Trial']==2][parameter].mean()-df.loc[df['Trial']==1][parameter].mean(),2)
    difft3t2=round(df.loc[df['Trial']==3][parameter].mean()-df.loc[df['Trial']==2][parameter].mean(),2)
    difft4t3=round(df.loc[df['Trial']==4][parameter].mean()-df.loc[df['Trial']==3][parameter].mean(),2)

    pvaluelist=result.iloc[:,3]
    symbollist=[]
    for p in pvaluelist:
        if p>0.01 and p<0.05:
            symbol='*'
        elif p<0.01:
            symbol='**'
        else:
            symbol='ns'
        symbollist.append(symbol)

    ax[i].annotate(str(difft2t1)+symbollist[0], xy=(0.19, 0.87), xytext=(0.19, 0.9), xycoords='axes fraction', 
                ha='center', va='bottom',
                arrowprops=dict(arrowstyle='-[, widthB=2, lengthB=0.1', lw=1.0),fontsize=20)
    ax[i].annotate(str(difft3t2)+symbollist[1], xy=(0.49, 0.87), xytext=(0.49, 0.9), xycoords='axes fraction', 
                ha='center', va='bottom',
                arrowprops=dict(arrowstyle='-[, widthB=3.5, lengthB=0.1', lw=1.0),fontsize=20)
    ax[i].annotate(str(difft4t3)+symbollist[2], xy=(0.8, 0.87), xytext=(0.8, 0.9), xycoords='axes fraction', 
                ha='center', va='bottom',
                arrowprops=dict(arrowstyle='-[, widthB=2, lengthB=0.1', lw=1.0),fontsize=20)

    ax[i].annotate('2h interval', xy=(0.2, 0.4), xytext=(0.2, 0.4), xycoords='axes fraction', 
                ha='center', va='bottom')
    ax[i].annotate('6dpf', xy=(0.2, 0.5), xytext=(0.2, 0.5), xycoords='axes fraction', 
                ha='center', va='bottom')
    ax[i].annotate('2h interval', xy=(0.8, 0.4), xytext=(0.8, 0.4), xycoords='axes fraction', 
                ha='center', va='bottom')
    ax[i].annotate('7dpf', xy=(0.8, 0.5), xytext=(0.8, 0.5), xycoords='axes fraction', 
                ha='center', va='bottom')
    ax[i].annotate('22h interval', xy=(0.5, 0.4), xytext=(0.5, 0.4), xycoords='axes fraction', 
                ha='center', va='bottom')
    ax[i].annotate('6dpf~7dpf', xy=(0.5, 0.5), xytext=(0.5, 0.5), xycoords='axes fraction', 
                ha='center', va='bottom')


    ax[i].set_ylabel(ylabels[i], fontsize=20)

plt.savefig('fig3C-E.pdf',bbox_inches = 'tight',
    pad_inches = 0)
plt.show()
```

In [25]:

```
figS2, ax = plt.subplots(1,2,figsize=(12,6))
plt.subplots_adjust(wspace=0.4)
ylabels=['LD-CI', 'PC-CI']
for i, par in enumerate (ylabels):
   
    df=df_pop_arena_trial.copy()
    df=df.loc[df['Trial']==1]

    t1=df.loc[(df.index.str.split('-').str[2].str.contains(r'(^[12][A-H])'))&(df.index.str.split('-').str[1].str.contains('Plate1'))][par]

    t2=df.loc[(df.index.str.split('-').str[2].str.contains(r'(^11|12[A-H])'))&(df.index.str.split('-').str[1].str.contains('Plate2'))][par]

    ax[i].violinplot([t1, t2], showmeans=True)
   
    pval=pg.ttest(t1, t2, paired=True, tail='two-sided').round(2).iloc[0,3]
    df=pg.ttest(t1, t2, paired=True, tail='two-sided').round(2).iloc[0,1]
    df=int(df)
    
    ax[i].spines['right'].set_visible(False)
    ax[i].spines['top'].set_visible(False)
    ax[i].set_xticks([1,2])
    ax[i].set_ylabel(ylabels[i],fontsize=20)
    ax[i].set_xticklabels(labels=['First group', 'Last group'],fontsize=20)
    ax[i].annotate('p = '+str(pval), xy=(0.4, 0.87), xytext=(0.4, 0.9), xycoords='axes fraction', fontsize=20)
    ax[i].annotate('df = '+str(df), xy=(0.4, 0.8), xytext=(0.4, 0.8), xycoords='axes fraction', fontsize=20)
plt.savefig('figS2.pdf',bbox_inches = 'tight',
    pad_inches = 0)   
plt.show()
```

```
/Users/jialexu/opt/anaconda3/lib/python3.7/site-packages/pandas/core/strings.py:1843: UserWarning: This pattern has match groups. To actually get the groups, use str.extract.
  return func(self, *args, **kwargs)
/Users/jialexu/opt/anaconda3/lib/python3.7/site-packages/pingouin/parametric.py:198: UserWarning: x and y have unequal sizes. Switching to paired == False. Check your data.
  warnings.warn("x and y have unequal sizes. Switching to "
```

In [26]:

```
from matplotlib.ticker import FormatStrFormatter
df=df_pop_arena_trial.copy().round(3)
def minmax(x):
    xmin=x.min()
    xmax=x.max()
    return (x-xmin)/(xmax-xmin)

def LTLI(tearly, tlate):

    return ((tlate/(0.5*(tearly+tlate))), (tearly/(0.5*(tearly+tlate))))
    
dic_6dpf={}
dic_7dpf={}
dic_overnight={}
Legendlabels=['LD-CI', 'PC-CI', 'No. of dark zone entry', 'Ave. duration of dark zone entry', 'Latency to 1st dark zone entry']
fig, ax=plt.subplots(1,3, figsize=(80,30), sharey=True,sharex=True)
plt.subplots_adjust(wspace=0.2)

for num, parameter in enumerate(df.columns[np.r_[3,4,5,6,7]]):
    t1=df.loc[df['Trial']==1][parameter]
    t1=minmax(t1)
    t2=df.loc[df['Trial']==2][parameter]
    t2=minmax(t2)
    t3=df.loc[df['Trial']==3][parameter]
    t3=minmax(t3)
    t4=df.loc[df['Trial']==4][parameter]
    t4=minmax(t4)
    if len(parameter.split('_'))==6:
        df_hab_6dpf=LTLI(t1, t2)[1]
        df_hab_7dpf=LTLI(t3, t4)[1]
        df_hab_overnight=LTLI(t2, t3)[1]
    else:
        df_hab_6dpf=LTLI(t1, t2)[0]
        df_hab_7dpf=LTLI(t3, t4)[0]
        df_hab_overnight=LTLI(t2, t3)[0]
    
    if parameter in ['LD-CI', 'PC-CI']:
        linewidth=10
    else:
        linewidth=2
    df_hab_6dpf=df_hab_6dpf.fillna(1)
    df_hab_6dpf[(df_hab_6dpf>0)&(df_hab_6dpf<2)&(df_hab_6dpf!=1)].plot(kind='kde', label=Legendlabels[num], ax=ax[0], linewidth=linewidth)
    ax[0].spines['top'].set_visible(False)
    ax[0].spines['right'].set_visible(False)
    ax[0].set_xticklabels(ax[0].get_xticks(), Fontsize=50)
    ax[0].set_xlabel('LTLI Trial 1 vs. 2 (2hr interval)', Fontsize=70)
    ax[0].set_yticklabels(ax[0].get_yticks(), Fontsize=50)
    ax[0].yaxis.set_major_formatter(FormatStrFormatter('%.1f'))
    ax[0].set_ylabel('Prob. Density', Fontsize=70)
    
    df_hab_7dpf[(df_hab_7dpf>0)&(df_hab_7dpf<2)&(df_hab_7dpf!=1)].plot(kind='kde', label=parameter, ax=ax[2], linewidth=linewidth)
    ax[2].spines['top'].set_visible(False)
    ax[2].spines['right'].set_visible(False)
    ax[2].set_xticklabels(ax[1].get_xticks(), Fontsize=50)
    ax[2].set_xlabel('LTLI Trial 3 vs. 4 (2hr interval)', Fontsize=70)
    
    df_hab_overnight[(df_hab_overnight>0)&(df_hab_overnight<2)&(df_hab_overnight!=1)].plot(kind='kde', label=parameter, ax=ax[1], linewidth=linewidth)  
    ax[1].spines['top'].set_visible(False)
    ax[1].spines['right'].set_visible(False)
    ax[1].set_xticklabels(ax[2].get_xticks(), Fontsize=50)
    ax[1].set_xlabel('LTLI Trial 2 vs. 3 (overnight interval)', Fontsize=70)  
    
    dic_6dpf.update({parameter:df_hab_6dpf})
    dic_7dpf.update({parameter:df_hab_7dpf})
    dic_overnight.update({parameter:df_hab_overnight})

handles, labels = ax[0].get_legend_handles_labels()
fig.legend(handles, labels, loc='upper center', 
           fontsize=60, ncol=len(Legendlabels),bbox_to_anchor=(0.45,0.06))
plt.subplots_adjust(wspace=0.2)
plt.savefig('fig4B.pdf',bbox_inches = 'tight',
    pad_inches = 0)
plt.show()
```

In [29]:

```
def LTHI_classify(LTHI, threshold):
    if LTHI<threshold:
        return 'S'
    elif LTHI>threshold:
        return 'H'
    elif LTHI==threshold:
        return 'NL'
    
df_hab_6dpf=pd.DataFrame(dic_6dpf)
df_hab_7dpf=pd.DataFrame(dic_7dpf)
df_hab_overnight=pd.DataFrame(dic_overnight)
df_hab_6dpf.fillna(1, inplace=True)
df_hab_7dpf.fillna(1, inplace=True)
df_hab_overnight.fillna(1,inplace=True)
df_LTHI_LDCI=pd.concat([df_hab_6dpf['LD-CI'],df_hab_7dpf['LD-CI'],df_hab_overnight['LD-CI']],axis=1)
df_LTHI_LDCI.columns=['LTHI_6dpf', 'LTHI_7dpf', 'Overnight']
df_LTHI_LDCI.insert(0, 'Category_6dpf', df_LTHI_LDCI['LTHI_6dpf'].apply(lambda x: LTHI_classify(x, 1)))
df_LTHI_LDCI.insert(1, 'Category_Overnight', df_LTHI_LDCI['Overnight'].apply(lambda x: LTHI_classify(x, 1)))
df_LTHI_LDCI.insert(2, 'Category_7dpf', df_LTHI_LDCI['LTHI_7dpf'].apply(lambda x: LTHI_classify(x, 1)))


fig4d4f, ax=plt.subplots(1,3, figsize=(35,35))
fig4d4f.subplots_adjust(wspace=0.25)
xlabels=['Trial 1 vs. 2 (2hr interval)','Trial 2 vs. 3 (overnight interval)','Trial 3 vs. 4 (2hr interval)']
for i, cat in enumerate (df_LTHI_LDCI.columns[:3]):

    df_LTHI_LDCI[cat].value_counts().reindex(['H', 'S', 'NL']).\
    plot.pie(autopct='%1.2f%%', ax=ax[i], explode=[0.02]*3,
             colors=['#ADD8E6','#FFA07A','#fe1f14'], fontsize=40)
    ax[i].set_ylabel('')
    ax[i].set_xlabel(xlabels[i], fontsize=45)
plt.savefig('fig4E-G.pdf',bbox_inches = 'tight',
    pad_inches = 0)
```

In [28]:

```
def LPCI_classify(LPCI, PCreshold):
    if LPCI<PCreshold:
        return 'S'
    if LPCI>PCreshold:
        return 'H'
    elif LPCI==PCreshold:
        return 'NL'
    
df_hab_6dpf=pd.DataFrame(dic_6dpf)
df_hab_7dpf=pd.DataFrame(dic_7dpf)
df_hab_overnight=pd.DataFrame(dic_overnight)
df_hab_6dpf.fillna(1, inplace=True)
df_hab_7dpf.fillna(1, inplace=True)
df_hab_overnight.fillna(1,inplace=True)
df_LTHI_PCCI=pd.concat([df_hab_6dpf['PC-CI'],df_hab_7dpf['PC-CI'],df_hab_overnight['PC-CI']],axis=1)
df_LTHI_PCCI.columns=['LPCI_6dpf', 'LPCI_7dpf', 'Overnight']
df_LTHI_PCCI.insert(0, 'Category_6dpf', df_LTHI_PCCI['LPCI_6dpf'].apply(lambda x: LPCI_classify(x, 1)))
df_LTHI_PCCI.insert(1, 'Category_Overnight', df_LTHI_PCCI['Overnight'].apply(lambda x: LPCI_classify(x, 1)))
df_LTHI_PCCI.insert(2, 'Category_7dpf', df_LTHI_PCCI['LPCI_7dpf'].apply(lambda x: LPCI_classify(x, 1)))


fig4d4f, ax=plt.subplots(1,3, figsize=(35,35))
fig4d4f.subplots_adjust(wspace=0.25)
xlabels=['Trial 1 vs. 2 (2hr interval)','Trial 2 vs. 3 (overnight interval)','Trial 3 vs. 4 (2hr interval)']
for i, cat in enumerate (df_LTHI_PCCI.columns[:3]):

    df_LTHI_PCCI[cat].value_counts().reindex(['H', 'S', 'NL']).\
    plot.pie(autopct='%1.2f%%', ax=ax[i], explode=[0.02]*3,
             colors=['#ADD8E6','#FFA07A','#fe1f14'], fontsize=40)
    ax[i].set_ylabel('')
    ax[i].set_xlabel(xlabels[i], fontsize=45)
plt.savefig('fig4H-J.pdf',bbox_inches = 'tight',
    pad_inches = 0)
```

In [107]:

```
##Transition rate of learning type of dark avoidance##

#Transition 1# 
dic1={}
for cat in ['H', 'S', 'NL']:
    summary=df_LTHI_LDCI.loc[df_LTHI_LDCI['Category_6dpf']==cat]['Category_Overnight'].value_counts().reindex(['H', 'S', 'NL'])
    summary=(100*summary/summary.sum()).round(2)
    dic1.update({cat:summary})
df1=pd.DataFrame(dic1)
df1.columns='6dpf_'+df1.columns
df1.index='Overnight_'+df1.index
df1.fillna(0, inplace=True)
df1.round(1)

#Transition 2#
dic2={}
for cat in ['H', 'S', 'NL']:
    summary=df_LTHI_LDCI.loc[df_LTHI_LDCI['Category_Overnight']==cat]['Category_7dpf'].value_counts().reindex(['H', 'S', 'NL'])
    summary=(100*summary/summary.sum()).round(2)
    dic2.update({cat:summary})
df2=pd.DataFrame(dic2)
df2.columns='Overnight_'+df2.columns
df2.index='7dpf_'+df2.index
df2.fillna(0, inplace=True)
df2.round(1)
```

Out[107]:

|  | Overnight\_H | Overnight\_S | Overnight\_NL |
| --- | --- | --- | --- |
| 7dpf\_H | 46.1 | 70.3 | 67.8 |
| 7dpf\_S | 53.9 | 25.6 | 0.0 |
| 7dpf\_NL | 0.0 | 4.1 | 32.2 |

In [108]:

```
##Transition rate of learning type of center avoidance##

#Transition 1#
dic1={}
for cat in ['H', 'S', 'NL']:
    summary=df_LTHI_PCCI.loc[df_LTHI_PCCI['Category_6dpf']==cat]['Category_Overnight'].value_counts().reindex(['H', 'S', 'NL'])
    summary=(100*summary/summary.sum()).round(2)
    dic1.update({cat:summary})
df1=pd.DataFrame(dic1)
df1.columns='6dpf_'+df1.columns
df1.index='Overnight_'+df1.index
df1.fillna(0, inplace=True)
df1.round(1)

#Transition 2#
dic2={}
for cat in ['H', 'S', 'NL']:
    summary=df_LTHI_PCCI.loc[df_LTHI_PCCI['Category_Overnight']==cat]['Category_7dpf'].value_counts().reindex(['H', 'S', 'NL'])
    summary=(100*summary/summary.sum()).round(2)
    dic2.update({cat:summary})
df2=pd.DataFrame(dic2)
df2.columns='Overnight_'+df2.columns
df2.index='7dpf_'+df2.index
df2.fillna(0, inplace=True)
df2.round(1)
```

Out[108]:

|  | Overnight\_H | Overnight\_S | Overnight\_NL |
| --- | --- | --- | --- |
| 7dpf\_H | 38.8 | 69.5 | 88.9 |
| 7dpf\_S | 61.2 | 29.5 | 0.0 |
| 7dpf\_NL | 0.0 | 1.0 | 11.1 |

In [31]:

```
##Power analysis with varied sample sizes##

#data analysis#
import random
samplesize=list(range(100,1700, 100))
dict1t2={}
dict2t3={}
dict3t4={}
for size in samplesize:
    resultlistt1t2=[]
    resultlistt2t3=[]
    resultlistt3t4=[]
    for i in range (0,100):    
        testlist=random.sample(list(df_pop_arena_trial.index.unique()), size)
        df=df_pop_arena_trial.loc[testlist]
        t1=df.loc[df['Trial']==1]['LD-CI']
        t2=df.loc[df['Trial']==2]['LD-CI']
        t3=df.loc[df['Trial']==3]['LD-CI']
        t4=df.loc[df['Trial']==4]['LD-CI']
        
        resultt1t2=pg.ttest(t1, t2, paired=True, tail='one-sided').round(2)
        resultt2t3=pg.ttest(t2, t3, paired=True, tail='one-sided').round(2)
        resultt3t4=pg.ttest(t3, t4, paired=True, tail='one-sided').round(2)
        
        resultlistt1t2.append(resultt1t2)
        resultlistt2t3.append(resultt2t3)
        resultlistt3t4.append(resultt3t4)
        
    valuet1t2=pd.concat(resultlistt1t2).iloc[:,-1]
    valuet2t3=pd.concat(resultlistt2t3).iloc[:,-1]
    valuet3t4=pd.concat(resultlistt3t4).iloc[:,-1]
    
    key=size
    dict1t2.update({key:valuet1t2})
    dict2t3.update({key:valuet2t3})
    dict3t4.update({key:valuet3t4})
    
dft1t2=pd.DataFrame(dict1t2)
dft2t3=pd.DataFrame(dict2t3)
dft3t4=pd.DataFrame(dict3t4)

#data visualization
#font ={'family' : 'Arial',
      #'size' : 11}

plt.figure(figsize=(12,5))
ax=dft1t2.mean().plot(marker='o',yerr=dft1t2.sem(), capsize=4, label='LD-CI changes between Trial 1 and Trial 2')
dft2t3.mean().plot(marker='D',yerr=dft1t2.sem(), capsize=4, label='LD-CI changes between Trial 2 and Trial 3', ax=ax)
dft3t4.mean().plot(marker='>',yerr=dft1t2.sem(), capsize=4, label='LD-CI changes between Trial 3 and Trial 4', ax=ax)
ax.spines['top'].set_visible(False)
ax.spines['right'].set_visible(False)

ax.legend()
ax.set_xticks(np.arange(100,1700,100))
ax.set_xlim(0,1800)
ax.set_xlabel('Population size')
ax.set_ylabel('Power')
plt.savefig('fig5.pdf',bbox_inches = 'tight',
    pad_inches = 0)
mpl.rc('font', **font)
plt.show()
```

In [ ]:
